# A Type VII-secreted toxin enables inter-mycobacterial competition

**DOI:** 10.64898/2026.03.26.714487

**Authors:** Samuel T. Benedict, Kieran Bowran, K.E. Eunice Lee, Jean-Lou Reyre, Cheng-Ruei Han, Huda Ahmad, Aaron Franklin, Nicole A. Mietrach, Abigail Layton, Emmanuele Severi, Kamilla Anochshenko, Gregory Goudge, Simon G. Caulton, Todd L. Lowary, Andrew L. Lovering, Manuel Banzhaf, Elisabeth C. Lowe, Tracy Palmer, Patrick J. Moynihan

## Abstract

Most bacteria live in complex environments where resources are scarce and competition is fierce. These organisms have evolved mechanisms to compete with other bacteria, often through the specialised secretion of proteinaceous toxins. Mycobacteria have not previously been reported to engage in this form of competition. The thick and unusual cell wall of mycobacteria, comprised of peptidoglycan, arabinogalactan and mycolic acids, is generally thought to be highly protective to these bacteria. Many enzymes have evolved to maintain this structure, including the GH183 family, which cleaves arabinogalactan. Here, we establish for the first time that some mycobacteria have weaponised a subset of these endo-D-arabinanases to enable inter-bacterial competition. We show that mycobacteria secrete an endo-D-arabinanase effector via the type VII secretion system that specifically targets the arabinogalactan layer of the Mycobacteriales cell envelope. Using structural biology and biochemistry, we identify the molecular basis for this activity and reveal a new protein family that protects the bacterium from the activity of this toxin. Finally, our data uncover widespread T7-secreted toxins in the Mycobacteriales, pointing to extensive inter-mycobacterial competition.

## Introduction

Mycobacterial species are often considered in isolation, likely due to the paradigm of *Mycobacterium tuberculosis,* which is typically thought of as existing in monoculture during infection. However, most mycobacteria are not pathogens and instead live in complex ecosystems. In these environments, it can be expected that they will need to compete for resources. This could include specialised mechanisms for metal or nutrient acquisition or through direct interference competition.

In other bacteria, this evolutionary pressure has driven the emergence of several mechanisms to deliver diverse anti-bacterial proteinaceous effectors. In Gram-negative organisms, competition is predominantly driven by the type VI secretion system, where toxic effectors are secreted into nearby cells through a contractile apparatus resembling a phage baseplate^1–3^. In Bacillota, proteinaceous effectors are targeted to bacterial cells via the type VIIb secretion system, which lacks a similar contractile apparatus^4–7^.

Up to five Type VIIa secretion systems (T7SSa) are encoded by mycobacteria, termed ESX-1-5. ESX-1 is an essential virulence factor in *M. tuberculosis,* and its inactivation is the key attenuating mutation in the *M. bovis* BCG vaccine strain^8^. The largest group of T7SSa substrates is the PE/PPE proteins, which are named for conserved motifs in their N-terminus and comprise 10% of the *M. tuberculosis* proteome^9^. These proteins heterodimerise through their N-termini to form helical bundles^10^. A conserved WxG motif in the PPE protein and a proximal YxxxD/E in the PE partner likely form a composite signal sequence^11,12^. PE/PPE proteins are associated with all ESX systems except ESX-4. Conversely, *M. tuberculosis* CpnT, the precursor of tuberculosis necrotising toxin, a host-targeting NAD^+^-glycohydrolase, requires ESX-4 for secretion^13,14^. The CpnT N-terminus is predicted to adopt a PE/PPE-like fold despite sharing little detectable sequence similarity.

The Mycobacteriales cell wall is unusual amongst bacteria in that it is a tripartite structure consisting of peptidoglycan, arabinogalactan and mycolic acids^15^. This structure is essential under most conditions and, combined with other cell envelope components, poses a substantial permeability barrier. We recently discovered the glycoside hydrolase family 183 (GH183), which likely enables arabinogalactan remodelling through cleavage of the D-arabinan backbone^16^. In this study, we demonstrate that some mycobacteria have weaponised a sub-family of these enzymes, which is T7-secretion dependent and mediates inter-mycobacterial competition.

### Mycobacteria encode a predicted type VII secreted GH183

Arabinogalactan forms a covalent bridge between the peptidoglycan and mycolic acid outer membrane and is essential for the viability of mycobacteria^17–19^. This polysaccharide is phylogenetically restricted to the Mycobacteriales, and D-arabinan is only known to be essential in this group of bacteria. The recently discovered GH183 endo-D-arabinofuranases specifically target the α-1,5 linkages within D-arabinan, and two of these housekeeping enzymes are found in virtually all mycobacteria^16^. During the work, leading to the identification of these enzymes, we recognised that some mycobacteria, including some prominent cystic fibrosis pathogen *Mycobacterium abscessus strains,* encoded an additional GH183 homolog (Fig. 1a).

**Fig. 1.**
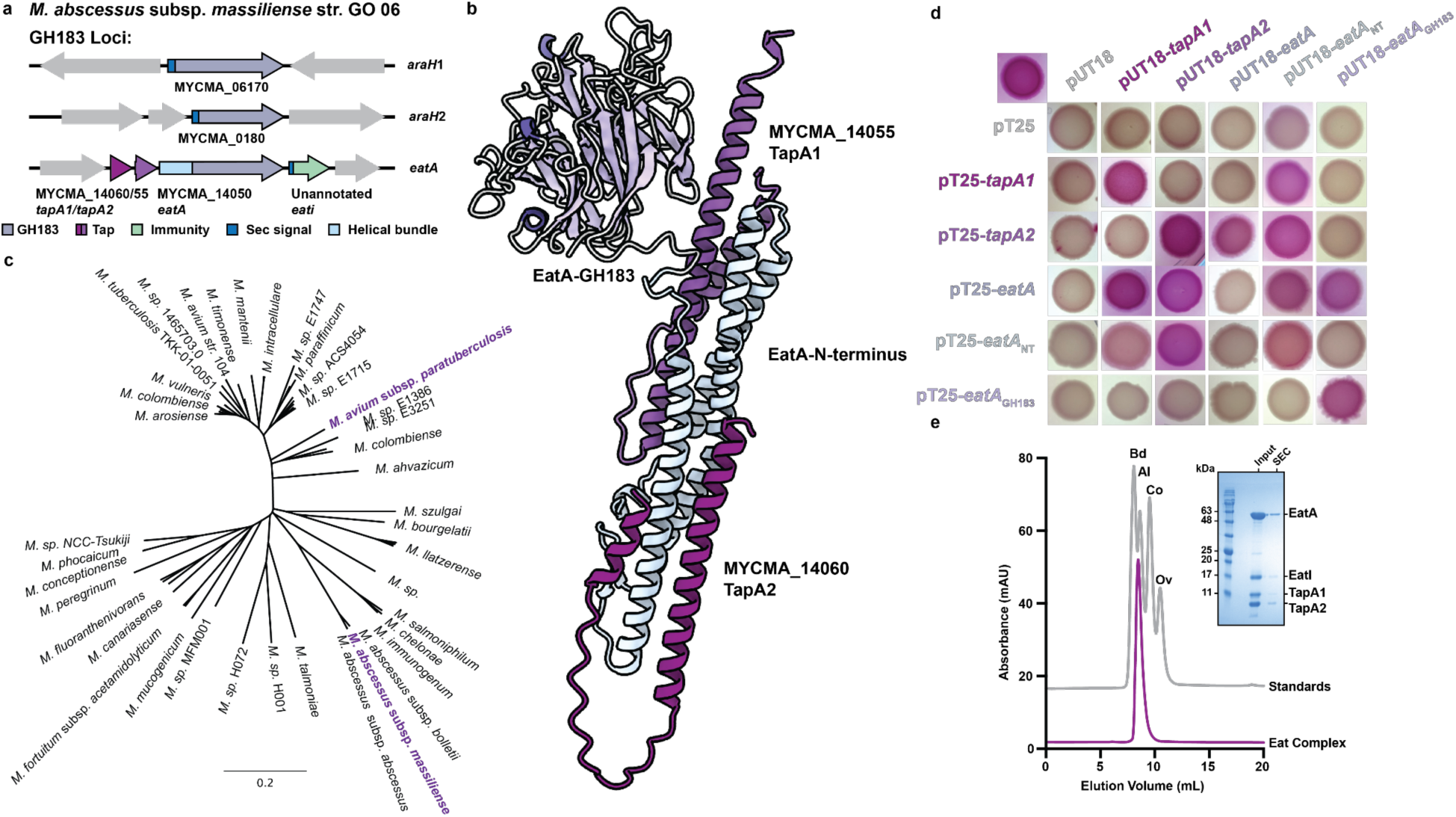
Mycobacteria encode a GH183 enzyme in a toxin-like locus. **a,** The *M. abscessus* subsp. *massiliense* str. GO 06 genome encodes two conserved housekeeping GH183 enzymes and a third copy with a locus structure including two small globular proteins encoded 5’ to the GH183 gene and a previously unassigned ORF downstream. AraH1, AraH2 and the unannotated ORF each have a predicted N-terminal Sec-secretion signal; the unannotated ORF also has a predicted lipo-box. **b,** The Alphafold3 prediction for TapA1 and TapA2 in complex with EatA reveals a predicted complex similar to other Type VII-secreted substrates. A copy of this prediction coloured by residue pLDDT is presented in Extended Data Fig. 2. **c,** An unrooted Foldtree using Foldseek score of cluster A0A7X5ZCJ1 demonstrates that the *eat* locus is distributed across the *Mycobacterium* genus in both slow-, and fast-growing organisms. Enzymes characterised here are shown in purple. **d,** Bacterial-two-hybrid results for *tap* genes and *eatA* illustrating interactions between the EatA N-terminus and both Tap proteins. **e,** Analytical size-exclusion of co-expressed genes from the *eat* locus demonstrating they elute as a single, high molecular weight complex. The input and eluted fractions were analysed by SDS-PAGE (inset), which showed the presence of all locus members in a single complex. Expected molecular weights: EatA - 56.9 kDa, EatI - 16.1 kDa, TapA1 - 12.7 kDa, TapA2 - 8.6 kDa, Complex - 94.3 kDa, blue dextran (Bd) - 200 kDa, aldolase (Al) - 158 kDa, conalbumin (Co) - 75 kDa, ovalbumin (Ov) - 44 kDa.

In *M. abscessus* subsp. *massiliense* str. GO 06 this protein, MYCMA_14050 is encoded at a locus with two small globular proteins (MYCMA_14055/60) and an unannotated open reading frame coding for a small lipoprotein (Fig. 1a). This locus organisation is reminiscent of the T7SSb toxin locus structure in Bacillota^4^. In Bacillota, T7SSb-secreted toxins usually have an N-terminal LxG domain, and they are co-encoded with helical partner proteins, termed **L**xG **a**ccessory **p**roteins (Lap) required for toxin secretion, and an immunity protein to prevent self-intoxication^5^. Given this locus organisation and the phylogenetic restriction of D-arabinan, we speculated that MYCMA_14050 may be involved in inter-mycobacterial competition. For reasons discussed later, we have renamed MYCMA_14050 as **E**sx-**a**ssociated **t**oxin A (EatA), with MYCMA_14055/60 designated TapA1/A2 (**T**oxin **a**ccessory **p**rotein), and the unannotated ORF as EatI (**Eat I**nhibitor).

AlphaFold 3 prediction of EatA revealed an elongated N-terminus rich in α-helices that interacts with TapA1 and TapA2 (Fig. 1b)^20^. Comparison of this prediction with structures of a T7SSa PE/PPE complex or toxin/accessory protein complexes from T7SSb demonstrates that the EatA complex includes appropriately positioned T7SS-targeting motifs (Extended Data Fig. 1,2a)^12,21^. Phylogenetic analysis of the EatA N-terminal region revealed it to cluster with helical domains from Bacillota T7SSb substrates rather than T7SSa substrates (Extended Data Fig. 1b). The presence of a GH183 domain and low sequence identity in the N-terminal helical domain of EatA complicated BLAST identification of orthologs. To circumvent this, we analysed the Foldseek cluster A0A7X5ZCJ1, to which EatA belongs, using Foldtree to determine the distribution of Eat (Fig. 1c)^22,23^. To our surprise, EatA orthologs are only found in mycobacterial species, with approximately 60 species possessing an EatA-like protein. This included both fast- and slow-growing environmental and pathogenic mycobacteria, at least one *Mycobacterium tuberculosis* strain and several uncultured isolates (Fig. 1c).

**Fig. 2.**
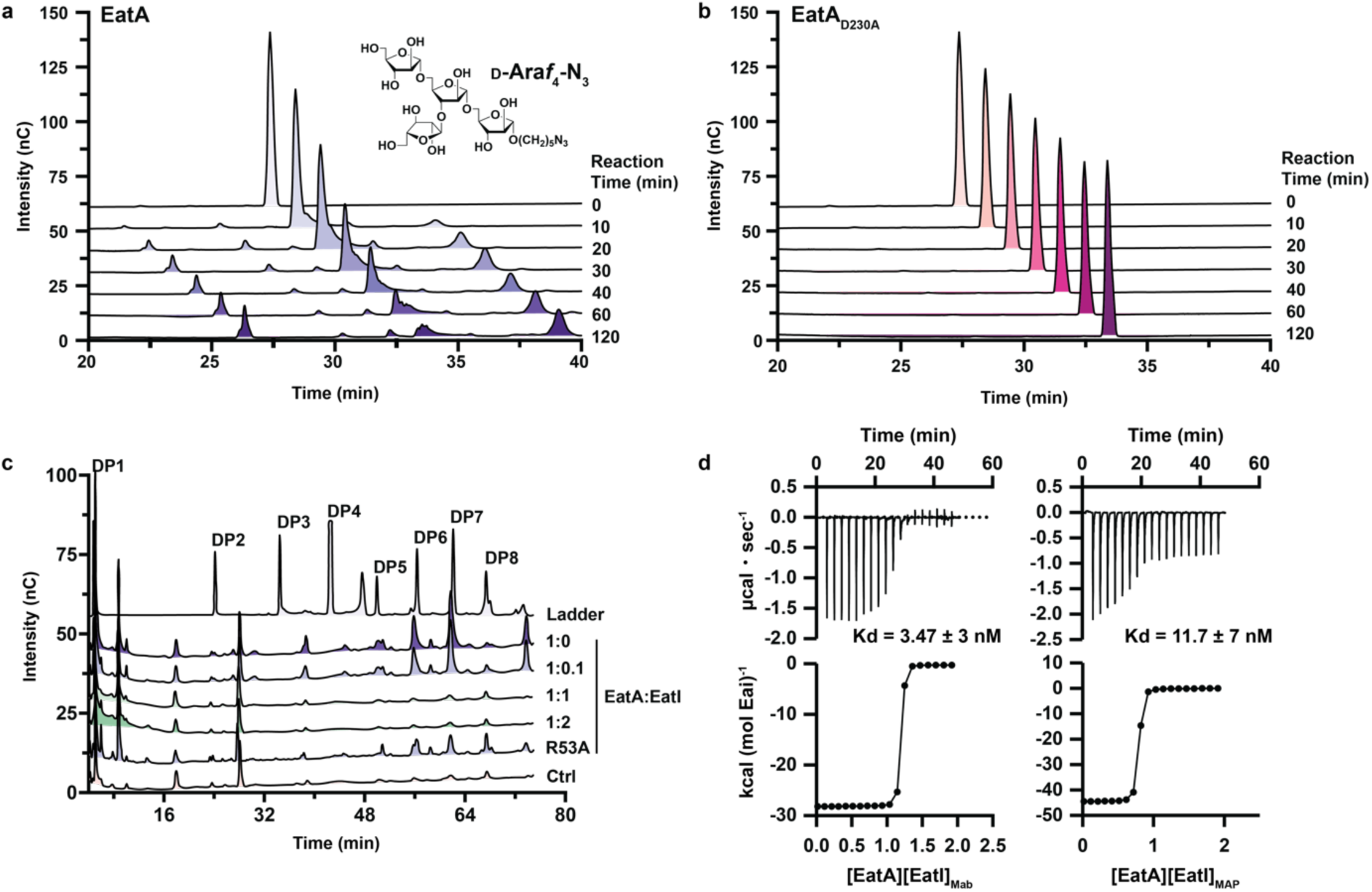
EatA proteins are GH183 family members and are inhibited by EatI. EatA_Mab_ (**a**) or EatA_MabD230A_ (**b**) (1 μM) were incubated with 50 µM D-Ara*f*_4_-N_3_ (structure inset) and at the indicated timepoints reactions were analysed by IC-PAD. Representative chromatograms of three biological replicates have been offset in the x-axis to aid visualisation. **c,** EatA_Mab_ (1 μM) was incubated either on its own or with the indicated molar ratios of EatI_Mab_ and 1 mg・mL^-1^ arabinogalactan from *M. smegmatis* mc^2^155. For EatI_MabR53A_, a 1:1 molar ratio was used. Representative chromatograms of three biological replicates are presented. The ladder is comprised of L-arabinofuranose oligosaccharides with the indicated degree of polymerisation (DP) and the signal has been scaled by half to aid visualisation. **d,** Isotherms of binding of EatA proteins to their cognate EatI proteins. Data is representative of two biological replicates. The top panel shows raw titration thermal changes in response to ligand addition, whilst the bottom panels show integrated peaks. Calculated Kd is shown between each panel.

### TapA1 and TapA2 form a complex with EatAMab

Because T7 secreted toxins from the Bacillota usually form complexes with small helical partners, which are then secreted^21,24^, we used a bacterial-two-hybrid assay with all cognate pairs of EatA, EatA_GH183_ (amino acids 185–526), EatA_N-term_ (amino acids 2–180), and TapA1/TapA2 to probe interactions (Fig. 1d). This revealed TapA1 and TapA2 to interact with both full-length EatA and its N-terminal domain, but not with each other, consistent with the Alphafold3 prediction (Fig. 1b,d)^20^. To further confirm this, we expressed the entire locus in *E. coli* isolating complexes via a His_6_ tag on EatA_Mab_ and a twin Strep-tag on TapA1. This resulted in a high-molecular-weight complex as determined by size exclusion chromatography, which contained all the proteins encoded at the locus (Fig. 1e).

### The Eat locus encodes an inhibitory protein

Glycoside hydrolases from the same families can catalyse very different activities, and so to confirm the catalytic function of EatA, we expressed and purified the GH183 domain from *M. abscessus* (AA 183-356). We then conducted hydrolysis assays using arabinogalactan and a synthetic D-arabinan oligosaccharide (D-Ara*f*_4_-N_3_; Fig. 2; Extended Data Fig. 4a,b). This confirmed the identity of EatA as an endo-D-arabinofuranase. Using the synthetic substrate, the enzyme had a specific activity of 0.027 ± 0.003 U・mg^-1^ (Extended Data Fig. 4b) and we observed no product formation when the catalytic acid/base was replaced with alanine (D230A) (Fig. 2b). The enzyme also displayed activity against arabinogalactan purified from *Mycobacterium smegmatis* mc^2^155 (Fig. 2c).

EatI is predicted to be a lipoprotein with no previously assigned function. We purified the *M. abscessus* protein without its predicted signal sequence (amino acids 25–172) and found that at an approximately 1:1 molar ratio of EatA_Mab_:EatI_Mab_, EatA activity was completely inhibited (Fig. 2c). Given its predicted localisation to the cell’s surface, we speculated that EatI binds directly to EatA, and so we combined the proteins and analysed them by size-exclusion chromatography (Extended Data Fig. 3). This revealed formation of a higher molecular weight complex, which we determined had a Kd of 3.47 ± 3 nM and a 1:1 molar ratio by isothermal titration calorimetry (Fig. 2d).

Given the conservation of this locus in many Mycobacteriales members, we purified the EatA GH183 domain (MAP44135_3721; Eat_MAP_; AA 214–540) and EatI (MAP44135_3722; Eai_MAP_; AA 22–170) from *Mycobacterium avium* subsp. *paratuberculosis* strain DSM 44135, which is a slow-growing mycobacterial species and the causative agent of Johne’s disease in ruminants. We confirmed similar biochemical functions for these proteins (Fig. 2b), with EatI_MAP_ inhibiting EatA_MAP_ (Fig. 2d) with a Kd of 11.7 ± 7 nM (Fig. 2d). We also found that the predicted immunity proteins did not form complexes and were not inhibitory for non-cognate pairs of *M. abscessus* and *M. avium* subsp. *paratuberculosis* proteins (Extended Data Fig. 3c, 4c,d).

### Molecular basis of Eat inhibition

To understand the molecular basis of EatA activity, we solved a 2.06 Å apo structure of the GH183 domain of EatA_Mab_ (A.A. 181-524; Fig. 3a). There were two monomers in the asymmetric unit, making a dimer via non-crystallographic symmetry which is unlikely to be physiologically relevant. Overall, each monomer resembles a five-bladed β-propeller. The two monomers are bound to a total of 11 ions that have been modelled as potassium, which are derived from the crystallisation condition. Similarly, the proteins are bound to several ethylene glycol molecules, with two bound in each active site (Fig. 3a, Extended Data Fig. 5a). The protein is structurally similar to the GH domain of EndoMA1 (PDB: 8IC1; Extended Data Fig. 5b), a characterised GH183 from *Microbacterium arabinogalactanolyticum* with an overall RMSD of 5.48 Å^25,26^. There are equivalent substrate-binding residues in EatA to most of those coordinating a modified tetra-D-arabinofuranoside ligand in EndoMA1 (Extended Data Fig. 5c). However, the loop containing P141 in EndoMA1 is significantly truncated in EatA, leading to a more open active site. This alignment supports the designation of D228 as the catalytic acid/base and D206 as the nucleophile in EatA, with an ethylene glycol molecule occupying approximately the same position as the -1 and -3 sugar-binding subsites (Extended Data Fig. 5c). The hydroxyl groups of these ethylene glycol molecules make several hydrogen bond interactions with residues that are conserved in both EndoMA1 and EatA, suggesting similar substrate specificity as supported by our biochemical assays (Fig. 2, Extended Data Fig. 4c). We next solved a 1.7 Å structure of the GH183 domain of EatA_Mab_ in complex with EatI_Mab_. Two complexes were observed in the asymmetric unit, which are nearly identical in structure (RMSD of 0.36 Å). The overall complex supports our size exclusion and isothermal titration calorimetry data, demonstrating a 1:1 stoichiometry for the GH183 and immunity domains, resulting in a buried surface area of 1628.5 Å^2^ ^27^. The immunity protein is comprised of a 10-strand β-sandwich made up of two β-meanders (Fig. 3b,d). To our knowledge, this is the first experimentally determined structure for this fold. Compared to the EatA_Mab_ apo structure, the binding of EatI_Mab_ induces a few conformational changes. Most notably, the small helical turn in EatA_Mab_ formed by P439−N441 becomes unwound, shifting the Cɣ of P439 by 7.4 Å towards EatI_Mab_ to make hydrophobic interactions with V46, Y60 and Y125. This positions D438 to form a salt bridge with H124 of EatI_Mab_ (Extended Data Fig. 5d). A second salt bridge is formed between R445 of EatA_Mab_ and E115 of EatI_Mab_. This rearrangement allows an extended β-hairpin of EatI_Mab_ to project approximately 22 Å into the active site of EatA_Mab_ placing the conserved R53 in the active site (Fig. 3b,d,f; Extended Data Fig. 5f). When compared to the EndoMA1 substrate complex (PDB 8IC1; Extended Data Fig. 5c), it becomes clear that the extended hairpin of EatI_Mab_ blocks the predicted -1 to -3 substrate-binding subsites, precluding catalytic activity (Fig. 3f). While structurally unrelated, this inhibitory mechanism is reminiscent of BliPI and II inhibition of TEM-1 and several families of lysozyme inhibitors^28–31^.

**Fig. 3.**
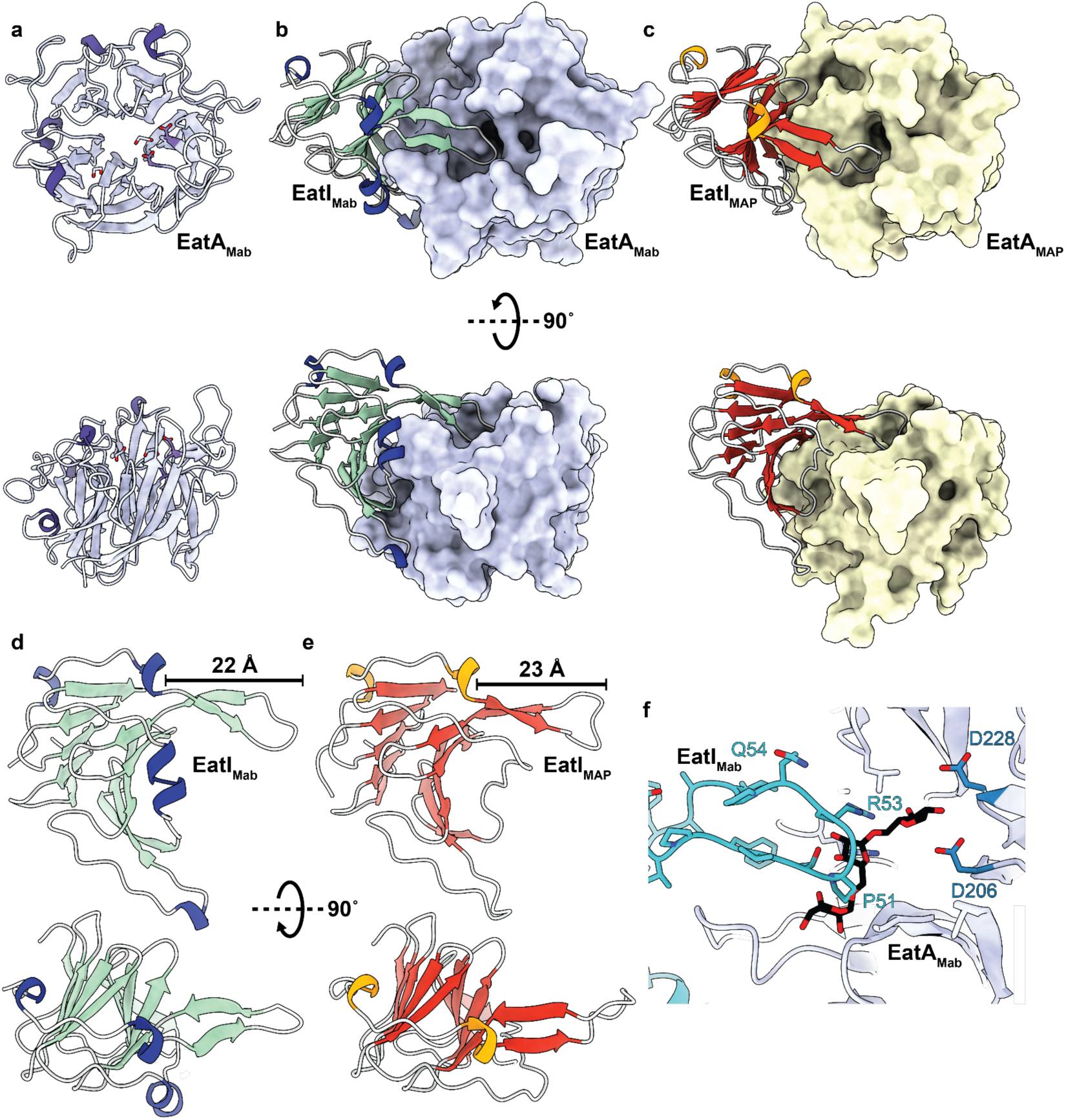
Molecular basis for EatI inhibition of EatA toxins. **a,** A 2.06 Å X-ray crystal structure of apo-EatA_Mab_ reveals a five-bladed β-propeller fold. **b,** The 1.7 Å structure of EatA-EatI_Mab_ reveals a 1:1 stoichiometry with the placement of an inhibitory loop from EatI_Mab_ in the active site of EatA_Mab,_ which is shown in the same pose as in panel **a** in lilac with surface representation. **c,** The 2.16 Å EatA-EatI_MAP_ complex is presented in the same orientation as the EatA-EatI_Mab_ structure, showing a similar overall fold and pose for both proteins. Structural alterations at the interface between these preclude cross-inhibition by non-cognate immunity proteins (Extended Data Fig. 5d,e). The overall fold of EatI from *M. abscessus* subsp. *massiliense* (**d,** green) and *M. avium* subsp. *paratuberculosis* (**e,** orange) is a β-sandwich with an elongated hairpin that forms the protein’s inhibitory loop. **f,** When aligned with the substrate (substrate shown in black) complex of EndoMA1 (PDB: 8IC1), it becomes evident that the inhibitory loop of EatI would preclude productive binding of the substrate.

To help understand why EatI proteins were not cross-protective *in vitro*, we also solved a 1.65 Å structure of the *M. avium* subsp. *paratuberculosis* EatA_MAP_–EatI_MAP_ complex (Fig. 3c,e). The structure is broadly similar to the EatA_Mab_–EatI_Mab_ complex (overall RMSD of 2.58 Å) with a buried interface of 1465.2 Å^2^. The binding of the EatI_MAP_ to EatA_MAP_ has several distinct features, however, likely explaining the lack of EatI cross-inhibition. EatA_MAP_ lacks the small interface helix, which is unwound in the EatA_Mab_–EatI_Mab_ complex. Instead, this loop is truncated by four residues to six amino acids in EatA_MAP_. Similarly, in EatI_Mab_, residues 119−124 form a helix absent in EatI_MAP_ (Fig. 3d,e, Extended Data Fig. 5d,e). This shifts the equivalent loop in EatI_MAP_ (amino acids 113−129) by 3.8 Å, which would clash with the unwound interface helix in EatA_Mab_. The net result of these structural changes is that the inhibitory β-hairpin in EatI_MAP_ is shifted by approximately 4 Å as it passes over the side of EatA_MAP_ relative to equivalent residues in the EatA–EatI_Mab_ complex (Extended Data Fig. 5e). Surprisingly, despite this shift, the arginine residue (R53/52) at the tip of the loop is placed in a nearly identical location in both structures indicating that the mechanism of substrate occlusion is likely the same for both complexes (Extended Data Fig. 5f). Supporting this, substituting R53 with alanine reduces EatI_Mab_ inhibition of EatA_Mab_ (Fig. 2c).

### EatA is toxic to mycobacteria via an ESX-4-dependent mechanism

*M. abscessus* ATCC 19977 lacks the *eatA* locus, and so to generate an experimentally tractable system, we constructed a strain that carried an L-arabinose inducible copy of the *M. abscessus* subsp. *massiliense* str. GO 06 *eatA* locus in the pMYBAD-K plasmid backbone^32^. Antagonism by the T7SS is contact-dependent in the Bacillota, and mycobacteria are non-motile^5^. We developed a pinning assay to measure mycobacterial fitness to overcome these limitations and investigate the potential for EatA-induced self-killing. We pinned the strain onto agar containing either repressor (D-glucose) or inducer (L-arabinose) and imaged the resulting colonies under standardised conditions. We used colony opacity as a proxy for cell fitness and observed a significant decrease in fitness in an *eatA-*dependent fashion (Fig. 4a). This decrease required a catalytically competent EatA and was rescued by including EatI (Fig. 4a). These data allow us to conclude that expression of EatA is toxic to *M. abscessus* and can be counteracted by expression of the cognate immunity protein.

**Fig. 4.**
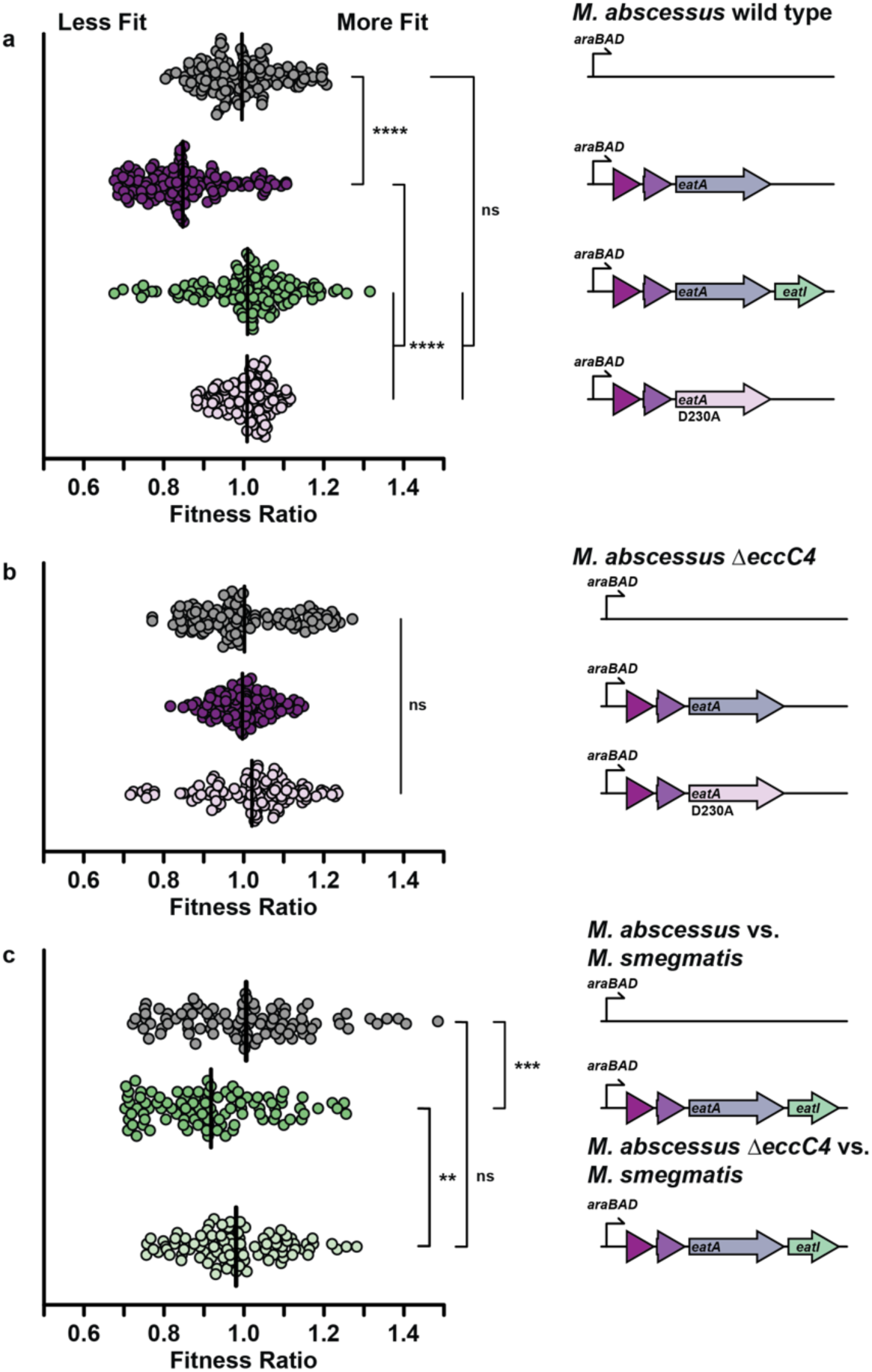
EatA proteins have toxic effects in mycobacteria. **a,** *M. abscessus* ATCC 19977 or **b,** *M. abscessus* ATCC 19977 Δ*eccC4* were transformed with a pMYBAD plasmid containing the indicated components of the *eatA* locus under the control of an arabinose inducible promoter. The D230A mutant of EatA is catalytically inactive. These were pinned (n = 128) onto tryptic soy agar with either 1% D-glucose or L-arabinose, grown for 48 h and imaged. Fitness is measured as a ratio of the colony opacity grown in non-inducing (D-glucose) or inducing (L-arabinose) conditions. **c,** The *M. abscessus* WT or Δ*eccC4* strain containing either an empty vector or the complete toxin locus was mixed at a 1:1 ratio with *M. smegmatis* mc^2^155 pTEC27-tdTomato. These were pinned (n = 96) onto solid media, and colony fluorescence was used as a proxy for *M. smegmatis* growth. Fitness is measured as the ratio of normalised colony fluorescence in inducing (L-arabinose) or non-inducing (D-glucose) conditions. Significance was determined using a one-way analysis of variance to compare differences from the control condition. **** = p < 0.0001, *** = p < 0.001, ** = p < 0.01.

ESX-4 T7SS is the ancestral T7SS system in mycobacteria, and its expression is induced by cell-cell contact^33,34^. We inactivated *M. abscessus* ATCC 19977 ESX-4 by deleting *eccC4,* which encodes part of the core T7SS apparatus^35–38^. Secretion of EsxU, a small *esx-4*-encoded effector, was abolished in the *eccC4* strain, confirming the inactivation of ESX-4 (Extended Data Fig. 6). Using the same colony pinning approach, we observed no *eatA*-dependent toxicity in the Δ*eccC4* strain, supporting our designation of secretion of this protein as Esx-4 dependent (Fig. 4b).

We next sought to determine if EatA could drive inter-species competition, so we made a prey strain by generating a strain of *M. smegmatis* mc^2^155 harbouring the pTEC27 plasmid, which includes a constitutively expressed tdTomato^39^. We combined this with the *M. abscessus* strain harbouring the complete *eat* locus at a 1:1 ratio and, as above, pinned them onto solid agar. To measure the fitness of the prey strain, we calculated the total fluorescence per colony when grown on L-arabinose or D-glucose and expressed this as a fitness ratio (Fig. 4c). Like the self-toxicity analysis above, we observed an EatA-dependent reduction in fitness for the prey strain (Fig. 4c). From this we conclude that EatA can drive inter-mycobacterial competition. These data demonstrate that mycobacteria produce a T7SS-dependent toxin that enables inter-mycobacterial competition.

### T7SS-dependent interbacterial toxins are broadly distributed in the Actinomycetota

Having established that mycobacteria utilise T7SS secretion for interbacterial competition, we sought to determine the diversity of toxic effectors in the Actinomycetota. Given the large number of available genomes and the varied environments in which it grows, we first analysed 946 *M. abscessus* strains available on NCBI. To determine the distribution of loci across *M. abscessus* and whether they followed the gene synteny described above, we queried our dataset for Tap-like proteins, yielding 1,439 unique protein sequences. Many of these were confirmed to be encoded in similar operons to the *eat* locus in discrete locations within the genome (Fig. 5a).

**Fig. 5.**
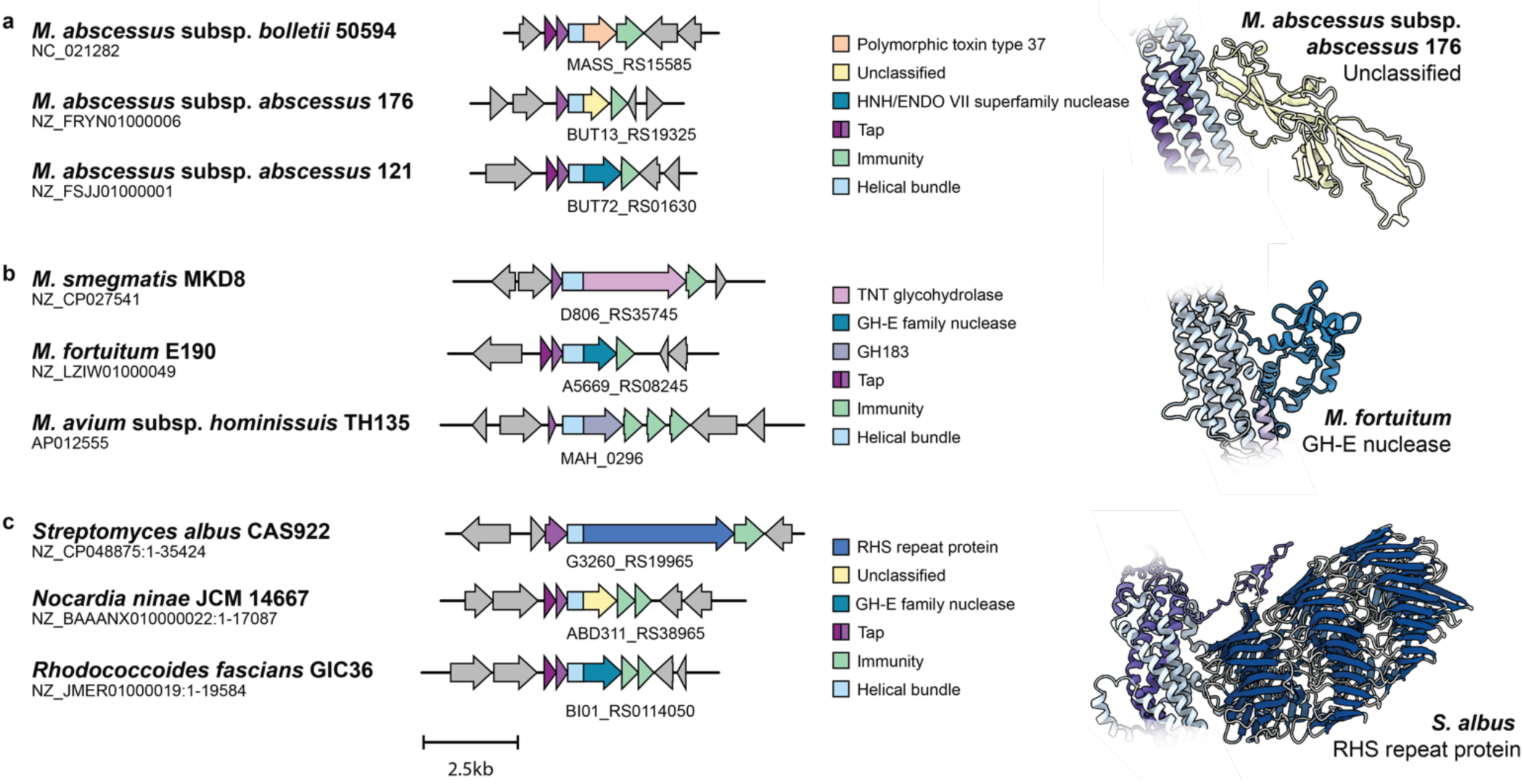
Actinomycetota bacteria encode diverse T7-secreted toxins. Representative *eat* loci from *M. abscessus* **a,** *Mycobacterium* **b,** and Actinomycetota **c,** strains. The helical bundle and upstream *tap*-like gene(s) are conserved features of these loci, which encode a diverse range of predicted toxin payloads (Extended Data Table 3). Genes are colour-coded for predicted function. The number of *tap*-like genes and predicted immunity proteins varies by locus. On the right are representative Alphafold 3 structural predictions for a toxin domain from each grouping.

These putative toxins span many predicted functions, including many known toxin families (Fig. 5a, Extended Data Table 3). The most commonly identified families were α/β hydrolase, nuclease HNH/ENDO superfamilies, NlpC/P60 family and TNT domain family (Extended Data Table 3). Proteins with no predicted function constituted 47% of all protein sequences. These proteins typically have between 400 and 1,000 residues, accompanied by one or two Lap/Tap-like encoding genes that mirror what was previously described for T7SSb substrates. Downstream-encoded immunity proteins tended to possess structural features reflecting the subcellular compartment of the corresponding proposed toxin target. For example, predicted extracellular targeting toxins would co-occur with adjacent immunities with predicted lipoprotein signal sequences or transmembrane helices.

Expanding this analysis to the Mycobacteriales highlights the presence of diverse Esx-dependent toxins within *Mycobacterium* and across the phylum (Fig. 5b,c), suggesting that inter-(myco)bacterial competition is a conserved feature.

## Discussion

In this study, we identify that mycobacteria use a T7-secreted proteinaceous effector to drive inter-mycobacterial competition. The EatA family targets cell wall arabinogalactan of the Mycobacteriales, which we have shown reduces fitness in a manner that requires catalytically active enzyme. Our identification of EatI proteins also shows how these bacteria protect themselves from self-killing, using a newly characterised protein fold as a specific and tight-binding EatA inhibitor. This killing mechanism is conceptually similar to the use of peptidoglycan-targeting effectors in other systems, such as Type VI-mediated interbacterial competition^40^.

The EatA system appears to be unusual amongst anti-bacterial effector proteins because it is biochemically restricted to a single phylum. Arabinogalactan is only found in the Mycobacteriales and is essential for their viability under most growth conditions. Indeed, its biosynthesis is the target of the front-line anti-tubercular drug ethambutol^41–45^. The existence of the *eatA* locus encouraged us to look for similar loci across mycobacteria, revealing numerous and diverse predicted T7-secreted toxic effectors. Excitingly, many of these loci encode predicted effector domains with no known function, pointing towards potential new targets for anti-mycobacterial therapy.

The competition for resources provides an enormous opportunity for evolution to shape bacterial genomes. The presence of ubiquitous T7-secreted toxin-mediated interference competition in the Mycobacteriales reframes our thinking of the evolutionary pressures placed on this phylum. *M. tuberculosis*, for example, is frequently thought of as living in isolation, and yet all strains of *M. tuberculosis* encode potential anti-bacterial T7-secreted toxins. Indeed, this points to a possible evolutionary origin for tuberculosis-necrotizing toxin, which is also ESX-4-dependent^14^.

Our study raises several important unresolved questions. For EatA to be active, the protein must access the periplasm of target cells. Many of the putative toxins we have identified are predicted to have cytoplasmic targets and correspondingly have predicted cytoplasmic immunity proteins. The mechanism by which these toxins transit the predator and prey cell envelopes remains unclear. A model by which the small Esx proteins facilitate tuberculosis-necrotizing toxin transport has been proposed; however, it is unclear how this would facilitate transport across three biological membranes and so would need to be validated in an inter-bacterial competition context^46^. It also remains to be determined if T7-secreted toxic effectors remain bound to their N-terminal carrier domain or if this is cleaved during secretion. Additionally, the mechanism by which toxin-secretion is regulated remains unclear and understanding this will be key to improving killing efficiency. Finally, the range of prey organisms that the T7 secretion system can target remains unknown but will significantly impact microbial ecology.

## Methods

### Strains and growth conditions

Unless stated otherwise, all chemicals and reagents were purchased from Sigma-Aldrich. *Escherichia coli* T7 Shuffle or BL21 DE3 (New England Biolabs) were used to express recombinant proteins, and *E. coli* 10-β was used for plasmid maintenance. *E. coli* strain M15 was used for over-production of the EatA-EatI-Tap protein complex. *M. smegmatis* mc^2^155 and *M. abscessus* ATCC 19977 were used in predation and self-toxicity experiments transformed through electroporation with the appropriate plasmids. *E. coli* strains, up to 100 mL, were grown in lysogeny broth. *E. coli* cultures used for protein purification were grown in terrific broth at 37 °C with agitation. *M. smegmatis* mc^2^155 and *M. abscessus* ATCC 19977 were grown in tryptic soy broth media for liquid cultivation and tryptic soy agar for solid growth at 37 °C. For mycobacteria, kanamycin and hygromycin selection were at 50 μg/mL.

### AlphaFold and FoldTree analysis

For protein structure prediction, amino acid sequences were inputted into the Alphafold 3 server and run using default settings^20^. Structural data were analysed using ChimeraX^25^. To identify EatA orthologs, the UniProt ID for EatA_Mab_ (A0A0U0ZL93) was input into the AlphaFold Clusters server, which identified Cluster A0A7X5ZCJ1^22^. This was then input into the Google Colab notebook for FoldTree, and the resulting tree was produced using FoldSeek score^23^. The resulting data were imported into Geneious Prime 2024.0.7 and are presented as an unrooted tree.

### Bacterial-2-Hybrid Assay

pT25 or pUT18 plasmid backbones encoding the appropriate bait or prey genes were transformed into *E. coli* BTH101^47^. These strains were spotted onto MacConkey agar supplemented with 1% D-maltose, 25 μg/mL chloramphenicol and 125 μg/mL ampicillin and grown at 30 °C until colonies were visible and control colonies were visibly pink before being photographed.

### Cloning

The expression plasmids for EatA_Mab_ (amino acids 183–356), EatI_Mab_ (amino acids 25– 172), EatA_MAP_ (amino acids 214–540), and EatI_MAP_ (amino acids 22–170) were synthesised and codon-optimised for *E. coli* expression by Twist Biosciences in a pET28a vector, including a hexahistidine tag and GSG linker on the N-terminus of EatA_Mab_, EatI_Mab_, and EatI_MAP_, proteins, and at the C-terminus of EatA_MAP_.

### Purification of the Eat Complex

A plasmid was constructed by ligating the *tapA1*/*A2*-*eatA*-*eatI* locus from *M. abscessus* M156, including a hexahistidine and twin-strep affinity tags to enable co-expression of the Eat complex, which we termed pqe70-*eatI*-*tapA1*-*tapA2*-TS-*eatA*-His_6_. This plasmid was transformed into *E. coli* M15 [pREP4] and was grown in 2 L of terrific broth (12 g/L tryptone, 24 g/L yeast extract, 4 mL/L glycerol, pH 7) to OD_600nm_ 0.6-1.0. Expression was induced using 1 mM IPTG for 4 h at 37 °C before harvesting and lysis in 50 mM HEPES pH 7.5, 150 mM NaCl, and 25 mM imidazole. Following lysis, we used a 3-step purification. This included a first purification using Ni-NTA resin in which the protein was bound and washed in the same buffer before batch elution with an increase to 500 mM imidazole. Fractions positive for the complex components were dialysed 100 mM Tris-HCl pH 8, 150 mM NaCl, 1 mM EDTA and loaded onto a Strep-TrapXT column. The protein was eluted in BXT buffer (IBA), concentrated, and dialysed into 25 mM HEPES pH 7.5, 150 mM NaCl before size-exclusion chromatography with a HiLoad 26/600 200 pg column.

### Purification of MAB and MAP EatA/EatI

*E. coli* T7 SHuffle (New England Biolabs) were transformed with 100 ng freshly prepped pET28a vector by heat shock and plated on Lysogeny Broth agar (VWR) plates supplemented with 50 μg/mL kanamycin. One plate of transformants was scraped into one litre of terrific broth before growing at 37 °C to an OD_600nm_ of 0.6-0.9 before cooling in cold water and inducing with 250 μM IPTG overnight at 16 °C. Cells were centrifuged at 5000 x *g* for 20 min before resuspending in a buffer containing 25 mM HEPES pH 8, 400 mM KCl, and 10 mM imidazole pH 8 with cOmplete protease inhibitor tablets (Roche). This cell suspension was lysed by sonication before clarifying the soluble fraction by centrifugation at 40,000 x *g* for 40 min at 4 °C. Crude proteins were isolated by immobilised metal affinity chromatography using cOmplete nickel resin (Roche). Fractions were assessed for purity by SDS-PAGE and then dialysed against 3 L of 25 mM HEPES pH 8, 200 mM KCl before further polishing by size exclusion chromatography using an Äkta Pure system using a HiLoad 26/600 200 pg column. Following polishing, proteins were concentrated using spin concentrators before snap freezing in liquid N_2_ and stored at -80 °C before use. Following freezing, protein concentrations were determined using the Pierce BCA protein assay kit (ThermoFisher Scientific) per the manufacturer’s instructions using the microplate protocol.

### Crystallography and data collection

For complex structures, toxin and immunity were mixed and equilibrated on ice for 20 min, ensuring a molar excess of immunity protein. Samples were then concentrated and polished again by size exclusion to remove excess immunity protein. The resulting purified complex was then concentrated. The crystallisation was conducted using the sitting drop vapour diffusion method at a 1:1 ratio with the crystallisation condition with a final volume of 2 μL. Sealed plates were incubated at 18 °C until crystal formation was observed.

For MAP_complex_, crystals formed in JCSG+ condition A10 (0.2 M ammonium nitrate pH 6.3, 20% PEG 3350) (Molecular Dimensions) in a 1:1 protein: precipitant ratio after approximately one week. Crystals possessed space group P212121 and were irradiated on beamline IO3 at Diamond Light Source at 100 K. For MAB_complex_, crystals formed Memgold2 condition C1 (0.075 M Magnesium chloride hexahydrate 0.1 M Sodium cacodylate 6.5 30 % w/v PEG 2000 MME) (Molecular Dimensions) in a 2:1 protein: precipitant ratio after approximately one week. Crystals formed in the space group P1211. For EatA_Mab_ apo, crystals formed in JCSG+ condition G10 (0.15 M potassium bromide, 30% PEG 3000-MME) (Molecular Dimensions) in a 1:1 protein: precipitant ratio after approximately one month. Crystals formed in the space group C2221 were irradiated on beamline IO3 at Diamond Light Source at 100 K.

All steps of phasing, solving, and structural refinement of datasets were carried out in the PHENIX suite (version 1.21rc1-5127-000). Phases were solved using PHASER using the appropriate Alphafold predicted structure and pruned to high confidence regions using the ‘Process Predicted Model’ package as a search model. Following phasing, successive rounds of manual and automated refinement using COOT (version 0.9.8.8) and PHENIX.refine, respectively, were carried out to build the final models.

### Isothermal Titration Calorimetry

Before analysis, aliquots of protein were dialysed for 3 h against 25 mM HEPES pH 8 with 200 mM KCl, to ensure identical buffers. EatA proteins were added to the chamber at 20 μM, and EatI was added to the syringe at 200 μM. Analyses were performed on a MicroCal PEAQ-ITC calorimeter at 25 °C using 19, 2 μL injections, with a reference power of 10 µcal. Data were analysed using MicroCal PEAQ-ITC Analysis Software (version 1.41).

### Analytical Size exclusion chromatography

Proteins were normalised to a concentration of 150 μM in 25 mM HEPES pH 8 and 200 mM NaCl. Chromatography was carried out at a flow rate of 0.8 mL/min on an Äkta pure system fitted with a Superdex 75 increase column, which had previously been equilibrated into the same buffer as above. The Eat and EatI proteins were mixed in a 1:1 molar ratio with either their cognate or non-cognate inhibitor proteins or buffer for single protein runs. Fractions were collected using the inline fraction collector and then subjected to 15% SDS-PAGE to analyse composition.

### Kinetic Analysis of EatA

To assess the degradation kinetics of EatA and EatA_D230A_ on D-Ara*f*_4_-N_3_, enzymes (1 µM) were incubated with 50 µM D-Ara*f*_4_-N_3_ at 37 °C in 50 mM citrate-phosphate buffer (pH 7). Reactions were conducted in technical triplicates and were terminated by adding an equal volume of 100 mM sodium hydroxide. The reaction products were analyzed by IC-PAD following the methods described above. A standard curve was generated to determine D-Ara*f*_4_-N_3_ concentrations, and data visualization was performed using GraphPad Prism 10.4.1.

### Ion Chromatography with Pulsed Amperometric Detection (IC-PAD)

The tetra-arabinofuranoside ligand (D-Ara*f*_4_-N_3_) used in kinetics investigations was synthesized using methods described previously^48,49^. The degradation kinetics of D-Ara*f*_4_-N_3_ and oligosaccharides derived from enzymatic arabinogalactan digestion were analysed using Ion Chromatography with Pulsed Amperometric Detection (IC-PAD). The analyses were performed on an ICS-6000 system equipped with a CARBOPAC PA-300 anion exchange column (ThermoFisher). Detection was facilitated by a gold working electrode and a PdH reference electrode, using a standard Carbo Quad waveform. Buffer A consisted of 100 mM NaOH, while Buffer B was composed of 100 mM NaOH and 0.5 M sodium acetate. For arabinogalactan product analysis, each sample was injected and run at a constant flow rate of 0.25 mL·min⁻¹ for 100 min using the following method: 0–10 min, isocratic elution with 100% Buffer A; 10–70 min, a linear gradient to 60% Buffer B; and 70–80 min, 100% Buffer B. The column was subsequently washed for 10 min with 500 mM NaOH, followed by re-equilibration for 10 min in 100% Buffer A.

For D-Ara*f*_4_-N_3_ analysis, the following method was applied: 0–5 min, isocratic elution with 100% Buffer A; 5–40 min, a linear gradient to 60% Buffer B; and 40–50 min, 100% Buffer B. The column was washed for 5 min with 500 mM NaOH, followed by re-equilibration in 100% Buffer A for 10 min. Commercially available L-arabinofuranoside-oligosaccharides (DP = 1–8) were used as standards (Megazyme) at a concentration of 25 µM. Data were processed using Chromeleon™ Chromatography Management System V.6.8, and final graphs were created using GraphPad Prism 10.4.1.

### EatA Inhibition Assays

Inhibition kinetics were assessed by incubating EatA (1 µM) and its cognate immunity protein at varying concentrations (0.1 µM, 1 µM, 2 µM) with arabinogalactan (1 mg·mL⁻¹) for 1 h at 37 °C in 50 mM citrate-phosphate buffer (pH 7). Prior to the reaction, EatA and the immunity protein were pre-incubated for 30 min at 4 °C in 50 mM citrate-phosphate buffer (pH 7). Reactions were conducted in technical triplicates and were terminated by adding an equal volume of 100 mM sodium hydroxide.

### Thin layer chromatographic enzyme assay

EatA and EatI proteins were normalised to 200 µM in 25 mM HEPES pH 8; 200 mM NaCl. EatA proteins were added to reactions at a final concentration of 5 µM. 1 mg/mL arabinogalactan purified from *M. smegmatis* mc^2^155 was used as the substrate, and 50 mM NaCl and 20 mM Tris-HCl (pH 8) were included in the reactions. Reactions were incubated for 1 h at 37 °C before boiling for 10 min to deactivate the enzyme. Ten μL of the reaction mixture was then spotted on silica gel TLC plates (Merck) and developed twice in 2:1:1 butan-1-ol: acetic acid: water before spraying with orcinol (5 g orcinol in 375 mL ethanol, 107 mL water, 16.2 mL concentrated sulfuric acid) and charred to reveal products.

### Pinning assays

For pinning competition assays, the prey strain was *M. smegmatis* mc^2^155 transformed with pTEC27, an L5-integrating plasmid constitutively expressing tdTomato^39^. Predator strains were *M. abscessus* ATCC 19977 transformed with pMYBADC-K with no insert or the *eat* genomic locus, including the two *tap* genes, *eatA* and *eatI* protein under the control of an arabinose inducible promoter or derivatives thereof. Mycobacterial cultures of predator and prey strains were grown in TSB with 0.05% Tween-80 with selection as appropriate to late stationary phase (OD_600_ = 2-4). Cultures were then pelleted at 3000 x *g* for 10 min before resuspending in the original volume of fresh TSB with 0.05% Tween-80. The culture was then diluted to an OD_600_ of 0.7 for *M. abscessus* self-intoxication assays and 0.5 for competition pinning assays. 100 µL per well of resulting cell suspension was transferred to a 96-well source plate (Sterilin) and pinned in a 384-spot format using the Biomatrix BM3-BC robot in single-well plates (VWR) onto either TSB or 7H9 plates solidified with 2% agar including 1% L-arabinose or D-glucose and incubated at 37 °C for 48 hours. Following incubation, the plates were imaged with an 18-megapixel Canon EOS Rebel T3i camera within the BM3-BC robot under controlled lighting. The S&P Imager (S&P Robotics, version 2.0.1.0) and EOS Utility (Canon, version 2.10.0.0) software programs were used for image processing.

### Colony Fitness Assay

Colony growth fitness was analysed using Iris software, quantifying colony opacity (version 0.9.7, available at https://github.com/critichu/Iris)^50^. Further analysis of Iris data was conducted using ChemGAPP small (available at https://github.com/HannahMDoherty/ChemGAPP/tree/main). Fitness scores were calculated relative to the non-induced condition. Bar plots of fitness ratios were generated by dividing the mean colony size by the mean colony size under a control condition, while swarm plots calculated fitness by dividing each colony size by the mean colony size under the control condition. Bar plots include 95% confidence intervals, and swarm plots display ANOVA significance levels.

### Fluorescence Data

Fluorescent data were collected using a BMG Polarstar plate reader. The excitation and emission wavelengths were 544 nm and 590 nm, respectively. Analysis was completed using the following R packages: imager, magick, ggplot2, reshape2, and ggbeeswarm^51–53^. The R script used for fluorescence data analysis is available at https://github.com/Hudaahmadd/Fluorescence-Analysis.

### Construction of Δ*eccC4* mutants in *M. abscessus*

Competent *M. abscessus* cells were electroporated with 500 ng of vector harbouring

1.5 kb regions flanking the target gene. Cells were recovered at 115 rpm at 37 °C for 4 hours and plated onto LB agar containing 50 μg/ml apramycin. Plates were incubated at 37 °C until the formation of pink colonies, followed by screening for genetic crossover events. Candidates were streaked onto LB agar containing 32 μg/ml isoniazid and incubated at 37 °C until the formation of white colonies. Candidates were patched onto either LB agar supplemented with apramycin or isoniazid to confirm resistance phenotypes and screened by PCR and whole genome sequencing to confirm the loss of the target gene.

### Design and purification of α-EccC4 antibody

Protein sequences of EccC_3_ (WP_005110740.1) and EccC_4_ (WP_005055990.1) were extracted from *M. abscessus* ATCC 19977 and aligned against EssC (Q932J9; residues 964 – 1479; *essC1* variant strain^54^ from *Staphylococcus aureus* Mu50 to identify the analogous cytoplasmic DIII domain in the mycobacterial proteins as previously described^55^. Plasmid construction and antibody production was carried out by GenScript. Briefly, mycobacterial genes were codon optimised and synthesised with the addition of an N-terminal hexahistidine-TEV fusion into pET28a(+), expressed within *E. coli* BL21 Star (DE3). Expression was induced with 0.5 mM IPTG for 16 h at 15 °C 200 rpm. Purified protein was obtained from the cell lysate supernatant by nickel affinity chromatography per manufacturer’s guidelines, with protein confirmed using monoclonal mouse-α-His (GenScript; Cat.No. A00186). Antibodies were raised in rabbits with three immunisation rounds, followed by ammonium sulfate precipitation.

### Purification and generation of α-EsxU antibodies

EsxU from *M. abscessus* was codon optimised and synthesised with the addition of an N-terminal hexahistidine-TEV fusion into pET28a(+) (GenScript) and subsequently transformed into *E. coli* BL21 (DE3). Cultures were grown until an OD_600_ of ∼0.6 – 0.8. Cells were induced with 0.5 mM IPTG for 4 hours at 37 °C, 200 rpm. Protein overproduction was confirmed via SDS-PAGE. Cells were harvested by centrifugation at 4,000 x g, with supernatants removed and pellets washed with 1X PBS followed by resuspension in Buffer A supplemented with cOmplete™ mini EDTA-free protease inhibitor cocktail (Sigma-Aldrich). Cells were lysed by sonication and pelleted by centrifugation, with the resultant supernatant passed through a 0.45 µm filter. The filtered supernatant was loaded onto an equilibrated HisTrap FF 5ml nickel Sepharose column (Cytiva), subsequently subjected to a denature wash with 50 mM Tris-HCl pH 7.5, 100 mM NaCl, 25 mM imidazole, 10% glycerol, 8M urea, washed in the same buffer lacking urea and eluted with the same buffer using a gradient of imidazole to 500 mM. Peak fractions were pooled, visualised by SDS-PAGE and concentrated using a 3 kDa spin concentrator (ThermoFisher Scientific). The concentrated samples were dialysed in 50 mM Tris-HCl pH 7.5, 100 mM NaCl and 10% glycerol overnight at 4 °C. Dialysed samples were incubated with 1 mg of TEV protease and left rolling at 4 °C. TEV protease-cleaved proteins were isolated through reapplication and reverse affinity nickel chromatography. Proteins were further purified by SEC with a Superdex 75 10/300 GL (Cytiva). Peak fractions were collected, concentrated and confirmed by SDS-PAGE.

Samples of EsxU were sent to Davids Biotechnologie to generate polyclonal antibodies. New Zealand rabbits were subjected to five protein immunisations over 63 days, followed by antigen-specific affinity purification. Antibodies were tested against purified protein samples to confirm activity and binding specificity.

### Analysis of putative T7SS-like proteins across Mycobacteriales

946 *M. abscessus* assemblies were downloaded from NCBI with an assembly level of scaffold or higher, packaged into local blastn (v2.13.0) and hmmer (v3.3.2) databases^56,57^. Both databases were queried under default search parameters using the WXG100 domains of known T7SSb substrates. Standard protein BLAST was used to query proteins across Mycobacteriales beyond *M. abscessus*. Candidate proteins were grouped based on C-terminal domain prediction by Motif Search using the Pfam database with default settings^58^. Topological features of corresponding immunity proteins were predicted through submission to SignalP 6.0^59^ and DeepTMHMM (v1.0.24)^60^. Genetic neighbourhoods were determined by submitting candidate proteins to the WebFlags^61^ server, with genetic variation assessed using both Cblaster^62^ and Clinker^63^ visualisation tools. Phylogenetic relationships were determined using MUSCLE alignment and constructing maximum likelihood trees with 250 bootstraps within MEGA (v10.1.8)^64,65^.

## Acknowledgements

This study was supported by the Wellcome Trust (through Investigator Award 224151/Z/21/Z to T.P. and 226644/Z/22/Z to E.C.L., T.P. and P.J.M. and a studentship to S.B.). A Newcastle University PhD studentship funded K.B. A PhD studentship funds E.L. through the Jeffcock & Luccock Endowments and the JJ Hunter Bequest. E.L. is the recipient of a Newcastle University Overseas Research Scholarship (NUORS). We thank Dr Eleanor Boardman for protein purification advice. The Biotechnology and Biosciences Research Council supported this work (grant BB/S010122/1 and BB/X00841X/1 to PJM, BB/X006298/1 to A.L.L. and BB/X016749/1 to ECL). K.A. is funded by a studentship from the Darwin Trust of Edinburgh. M.B. is funded through a UKRI Future Leaders Fellowship (MR/V027204/1) and a Springboard award (SBF005/1112).

## Competing interests

The authors declare no competing interests.

## Notes

### Competing Interest Statement

The authors have declared no competing interest.

## References

1. Pukatzki, S. et al. Identification of a conserved bacterial protein secretion system in Vibrio cholerae using the Dictyostelium host model system. Proc. Natl. Acad. Sci. 103, 1528–1533 (2006).

2. Basler, M., Pilhofer, M., Henderson, G. P., Jensen, G. J. & Mekalanos, J. J. Type VI secretion requires a dynamic contractile phage tail-like structure. Nature 483, 182–186 (2012).

3. Borgeaud, S., Metzger, L. C., Scrignari, T. & Blokesch, M. The type VI secretion system of Vibrio cholerae fosters horizontal gene transfer. Science 347, 63–67 (2015).

4. Cao, Z., Casabona, M. G., Kneuper, H., Chalmers, J. D. & Palmer, T. The type VII secretion system of Staphylococcus aureus secretes a nuclease toxin that targets competitor bacteria. Nat Microbiol 2, 16183 (2016).

5. Whitney, J. C. et al. A broadly distributed toxin family mediates contact-dependent antagonism between gram-positive bacteria. eLife 6, e26938 (2017).

6. Garrett, S. R. et al. A type VII-secreted lipase toxin with reverse domain arrangement. Nat. Commun. 14, 8438 (2023).

7. Ulhuq, F. R. et al. A membrane-depolarizing toxin substrate of the Staphylococcus aureus type VII secretion system mediates intraspecies competition. Proc. Natl. Acad. Sci. 117, 20836–20847 (2020).

8. Lewis, K. N. et al. Deletion of RD1 from Mycobacterium tuberculosis Mimics Bacille Calmette-Guérin Attenuation. J. Infect. Dis. 187, 117–123 (2003).

9. Cole, S. T. et al. Deciphering the biology of Mycobacterium tuberculosis from the complete genome sequence. Nature 393, 537–544 (1998).

10. Strong, M. et al. Toward the structural genomics of complexes: Crystal structure of a PE/PPE protein complex from Mycobacterium tuberculosis. Proc. Natl. Acad. Sci. 103, 8060–8065 (2006).

11. Daleke, M. H. et al. General secretion signal for the mycobacterial type VII secretion pathway. Proc. Natl. Acad. Sci. 109, 11342–11347 (2012).

12. Korotkova, N. et al. Structure of the Mycobacterium tuberculosis type VII secretion system chaperone EspG5 in complex with PE25–PPE41 dimer. Mol. Microbiol. 94, 367–382 (2014).

13. Lafuente, B. I., Ummels, R., Kuijl, C., Bitter, W. & Speer, A. Mycobacterium tuberculosis Toxin CpnT Is an ESX-5 Substrate and Requires Three Type VII Secretion Systems for Intracellular Secretion. mBio 12, 10.1128/mbio.02983-20 (2021).

14. Sun, J. et al. The Tuberculosis Necrotizing Toxin kills macrophages by hydrolyzing NAD. Nat Struct Mol Biol 22, 672–678 (2015).

15. Dulberger, C. L., Rubin, E. J. & Boutte, C. C. The mycobacterial cell envelope — a moving target. Nat Rev Microbiol 18, 47–59 (2020).

16. Al-Jourani, O. et al. Identification of d-arabinan-degrading enzymes in mycobacteria. Nat. Commun. 14, 2233 (2023).

17. Takayama, K. & Kilburn, J. O. Inhibition of synthesis of arabinogalactan by ethambutol in Mycobacterium smegmatis. Antimicrob Agents Ch 33, 1493–1499 (1989).

18. Mikusová, K., Slayden, R. A., Besra, G. S. & Brennan, P. J. Biogenesis of the mycobacterial cell wall and the site of action of ethambutol. Antimicrob. Agents Chemother. 39, 2484–2489 (1995).

19. Belanger, A. E. et al. The embAB genes of Mycobacterium avium encode an arabinosyl transferase involved in cell wall arabinan biosynthesis that is the target for the antimycobacterial drug ethambutol. Proc National Acad Sci 93, 11919–11924 (1996).

20. Abramson, J. et al. Accurate structure prediction of biomolecular interactions with AlphaFold 3. Nature 630, 493–500 (2024).

21. Klein, T. A. et al. Structure of a tripartite protein complex that targets toxins to the type VII secretion system. Proc. Natl. Acad. Sci. 121, e2312455121 (2024).

22. Barrio-Hernandez, I. et al. Clustering predicted structures at the scale of the known protein universe. Nature 622, 637–645 (2023).

23. 23.Moi, D. et al. Structural phylogenetics unravels the evolutionary diversification of communication systems in gram-positive bacteria and their viruses. bioRxiv 2023.09.19.558401 (2023) doi:10.1101/2023.09.19.558401.

24. Yang, Y. et al. Three small partner proteins facilitate the type VII-dependent secretion of an antibacterial nuclease. mBio 14, e02100–23 (2023).

25. Meng, E. C. et al. UCSF ChimeraX: Tools for structure building and analysis. Protein Sci. 32, e4792 (2023).

26. Shimokawa, M. et al. Identification and characterization of endo-α-, exo-α-, and exo-β-d-arabinofuranosidases degrading lipoarabinomannan and arabinogalactan of mycobacteria. Nat. Commun. 14, 5803 (2023).

27. Krissinel, E. & Henrick, K. Inference of Macromolecular Assemblies from Crystalline State. J. Mol. Biol. 372, 774–797 (2007).

28. Lim, D. et al. Crystal structure and kinetic analysis of β-lactamase inhibitor protein-II in complex with TEM-1 β-lactamase. Nat Struct Biol 8, 848–852 (2001).

29. Abergel, C. et al. Structure and evolution of the Ivy protein family, unexpected lysozyme inhibitors in Gram-negative bacteria. Proc. Natl. Acad. Sci. 104, 6394–6399 (2007).

30. Yum, S. et al. Structural basis for the recognition of lysozyme by MliC, a periplasmic lysozyme inhibitor in Gram-negative bacteria. Biochem. Biophys. Res. Commun. 378, 244– 248 (2009).

31. Leysen, S. et al. The structure of the proteinaceous inhibitor PliI from Aeromonas hydrophila in complex with its target lysozyme. Acta Crystallogr. Sect. D: Biol. Crystallogr. 71, 344–351 (2015).

32. Beckham, K. S. H., Staack, S., Wilmanns, M. & Parret, A. H. A. The pMy vector series: A versatile cloning platform for the recombinant production of mycobacterial proteins in Mycobacterium smegmatis. Protein Sci Publ Protein Soc 29, 2528–2537 (2020).

33. Clark, R. R., et al. Direct cell–cell contact activates SigM to express the ESX-4 secretion system in Mycobacterium smegmatis. Proc. Natl. Acad. Sci. 115, E6595–E6603 (2018).

34. Newton-Foot, M., Warren, R. M., Sampson, S. L., Helden, P. D. van & Pittius, N. C. G. van. The plasmid-mediated evolution of the mycobacterial ESX (Type VII) secretion systems. BMC Evol. Biol. 16, 62 (2016).

35. Beckham, K. S. H. et al. Structure of the mycobacterial ESX-5 type VII secretion system membrane complex by single-particle analysis. Nat. Microbiol. 2, 17047 (2017).

36. Beckham, K. S. H. et al. Structure of the mycobacterial ESX-5 type VII secretion system pore complex. Sci. Adv. 7, eabg9923 (2021).

37. Famelis, N. et al. Architecture of the mycobacterial type VII secretion system. Nature 576, 321–325 (2019).

38. Poweleit, N. et al. The structure of the endogenous ESX-3 secretion system. eLife 8, e52983 (2019).

39. Takaki, K., Davis, J. M., Winglee, K. & Ramakrishnan, L. Evaluation of the pathogenesis and treatment of Mycobacterium marinum infection in zebrafish. Nat. Protoc. 8, 1114–1124 (2013).

40. Russell, A. B. et al. Type VI secretion delivers bacteriolytic effectors to target cells. Nature 475, 343–347 (2011).

41. Safi, H., Sayers, B., Hazbón, M. H. & Alland, D. Transfer of embB Codon 306 Mutations into Clinical Mycobacterium tuberculosis Strains Alters Susceptibility to Ethambutol, Isoniazid, and Rifampin. Antimicrob. Agents Chemother. 52, 2027–2034 (2008).

42. Zhao, L. et al. Analysis of embCAB Mutations Associated with Ethambutol Resistance in Multidrug-Resistant Mycobacterium tuberculosis Isolates from China. Antimicrob. Agents Chemother. 59, 2045–2050 (2015).

43. Brossier, F. et al. Molecular Analysis of the embCAB Locus and embR Gene Involved in Ethambutol Resistance in Clinical Isolates of Mycobacterium tuberculosis in France. Antimicrob. Agents Chemother. 59, 4800–4808 (2015).

44. Sun, Q. et al. Mutations within embCAB Are Associated with Variable Level of Ethambutol Resistance in Mycobacterium tuberculosis Isolates from China. Antimicrob. Agents Chemother. 62, 10.1128/aac.01279-17 (2017).

45. Zhang, L. et al. Structures of cell wall arabinosyltransferases with the anti-tuberculosis drug ethambutol. Science 368, 1211–1219 (2020).

46. Tak, U., Dokland, T. & Niederweis, M. Pore-forming Esx proteins mediate toxin secretion by Mycobacterium tuberculosis. Nat. Commun. 12, 394 (2021).

47. Karimova, G., Pidoux, J., Ullmann, A. & Ladant, D. A bacterial two-hybrid system based on a reconstituted signal transduction pathway. Proc. Natl. Acad. Sci. 95, 5752–5756 (1998).

48. Ayers, J. D., Lowary, T. L., Morehouse, C. B. & Besra, G. S. Synthetic arabinofuranosyl oligosaccharides as Mycobacterial arabinosyltransferase substrates. Bioorg Med Chem Lett 8, 437–442 (1998).

49. Han, J., Gadikota, R. R., McCarren, P. R. & Lowary, T. L. Synthesis of octyl arabinofuranosides as substrates for mycobacterial arabinosyltransferases. Carbohydr. Res. 338, 581–588 (2003).

50. Kritikos, G. et al. A tool named Iris for versatile high-throughput phenotyping in microorganisms. Nat Microbiol 2, 17014–17014 (2017).

51. Ooms, J. Magick 1.0: Advanced Graphics and Image Processing in R. (2017) doi:10.59350/th49p-22b65.

52. Wickham, H., Chang, W. & Wickham, M. H. Package ‘ggplot2.’ Create elegant data visualisations using the grammar of graphics. Version 2, 1–189 (2016).

53. Wickham, H. Reshaping Data with the reshape Package. J. Stat. Softw. 21, (2007).

54. Warne, B. et al. The Ess/Type VII secretion system of Staphylococcus aureus shows unexpected genetic diversity. BMC Genom. 17, 222 (2016).

55. Kneuper, H. et al. Heterogeneity in ess transcriptional organization and variable contribution of the Ess/Type VII protein secretion system to virulence across closely related Staphylocccus aureus strains. Mol. Microbiol. 93, 928–943 (2014).

56. Altschul, S. F., Gish, W., Miller, W., Myers, E. W. & Lipman, D. J. Basic local alignment search tool. J. Mol. Biol. 215, 403–410 (1990).

57. Eddy, S. R. Accelerated Profile HMM Searches. PLoS Comput. Biol. 7, e1002195 (2011).

58. Mistry, J. et al. Pfam: The protein families database in 2021. Nucleic Acids Res. 49, D412–D419 (2020).

59. Teufel, F. et al. SignalP 6.0 predicts all five types of signal peptides using protein language models. Nat Biotechnol 1–3 (2022) doi:10.1038/s41587-021-01156-3.

60. Hallgren, J. et al. DeepTMHMM predicts alpha and beta transmembrane proteins using deep neural networks. (2022) doi:10.1101/2022.04.08.487609.

61. Saha, C. K., Pires, R. S., Brolin, H., Delannoy, M. & Atkinson, G. C. FlaGs and webFlaGs: discovering novel biology through the analysis of gene neighbourhood conservation. Bioinformatics 37, 1312–1314 (2020).

62. Gilchrist, C. L. M., et al. cblaster: a remote search tool for rapid identification and visualization of homologous gene clusters. Bioinform. Adv. 1, vbab016 (2021).

63. Gilchrist, C. L. M. & Chooi, Y.-H. clinker & clustermap.js: automatic generation of gene cluster comparison figures. Bioinformatics 37, 2473–2475 (2021).

64. Kumar, S., Stecher, G., Li, M., Knyaz, C. & Tamura, K. MEGA X: Molecular Evolutionary Genetics Analysis across Computing Platforms. Mol. Biol. Evol. 35, 1547–1549 (2018).

65. Stecher, G., Tamura, K. & Kumar, S. Molecular Evolutionary Genetics Analysis (MEGA) for macOS. Mol. Biol. Evol. 37, 1237–1239 (2020).

