## Extended Data for "A Type VII-secreted toxin enables inter-mycobacterial competition"

### Extended materials for: A Type VII-secreted toxin enables inter-mycobacterial competition

**Extended Data Fig. 1. Type VIIa and VIIb secretion motifs are conserved across diverse substrates.**

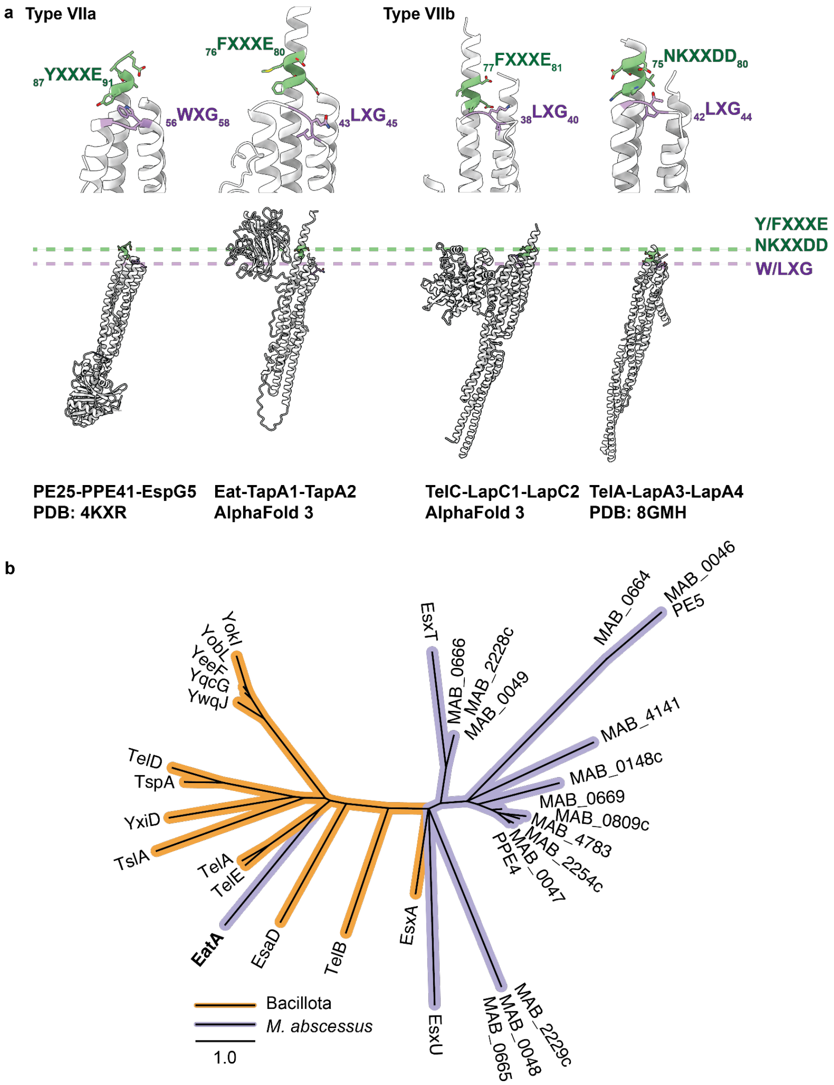


**a,** Across both Type VIIa and VIIb secretion systems, two motifs have been experimentally determined to be essential for cargo secretion. These are the W/LXG and Y/FXXXE (or NKXXDD) (upper panel). Alignment of the AlphaFold 3 or experimentally determined structures for EatA-TapA1-TapA2, TelC-LapC1-LapC2 and TelA-LapA3-LapA4 complexes to the experimentally determined structure of PE-25-PPE41-EspG5 demonstrates conservation of these motifs across Type VII substrates (lower panel). The dashed lines indicate the location of the motifs in the individual structures. AlphaFold statistics are presented in Extended Data Fig. 2. **b,** Phylogenetic analysis of the N-terminal helical domain of TVII substrates from the Bacillota (orange) and *M. abscessus* ATCC 19977. Protein sequences for all proteins annotated as PE or PPE and selected Bacillota T7b substrates were chosen for analysis. This was restricted to the N-terminal 200 (TelA-E, YxiD, TspA, YwQJ, YqcG, YeeF, YobL, YokL, EsaD, EatA, and MAB_4141, 0148, 0669, 0809c, 4783, 2254c, 0047, PPE4), 97 (EsxA, EsxT, MAB_2229c, 0048, 0665), 96 (MAB_0666, MAB_2228c), 102 (MAB_0664, 0046, PE5), 92 (EsxU) and C-terminal 199 (TslA) amino acids to capture the helical bundle domain.

**Extended Data Fig. 2. AlphaFold3 prediction of EatA-Tap1-Tap2 and TelC-LapC1-LapC2 complexes.**

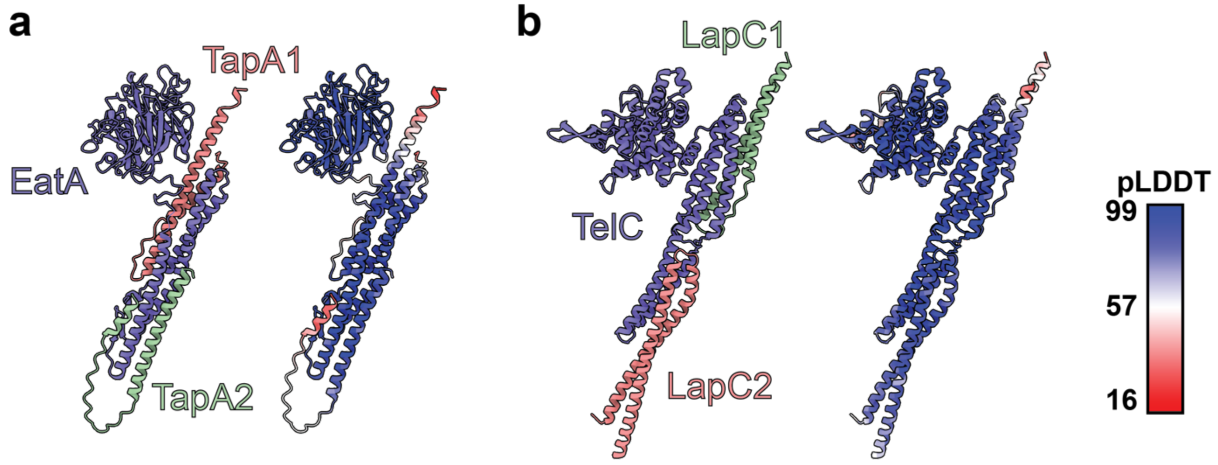


The amino acid sequences for **(a)** EatA, TapA1 and TapA2 (MYCMA_14050/60/55) and **(b)** TelC-LapC1-LapC2 (SIR_1489/90/91) were inputted to the AlphaFold 3 server using a single copy of each protein. For each protein, the predicted structure is shown on the left, coloured by chain, and on the right, according to per-atom pLDDT.

**Extended Data Fig. 3. Analytical size exclusion of EatA-EatI complexes.**

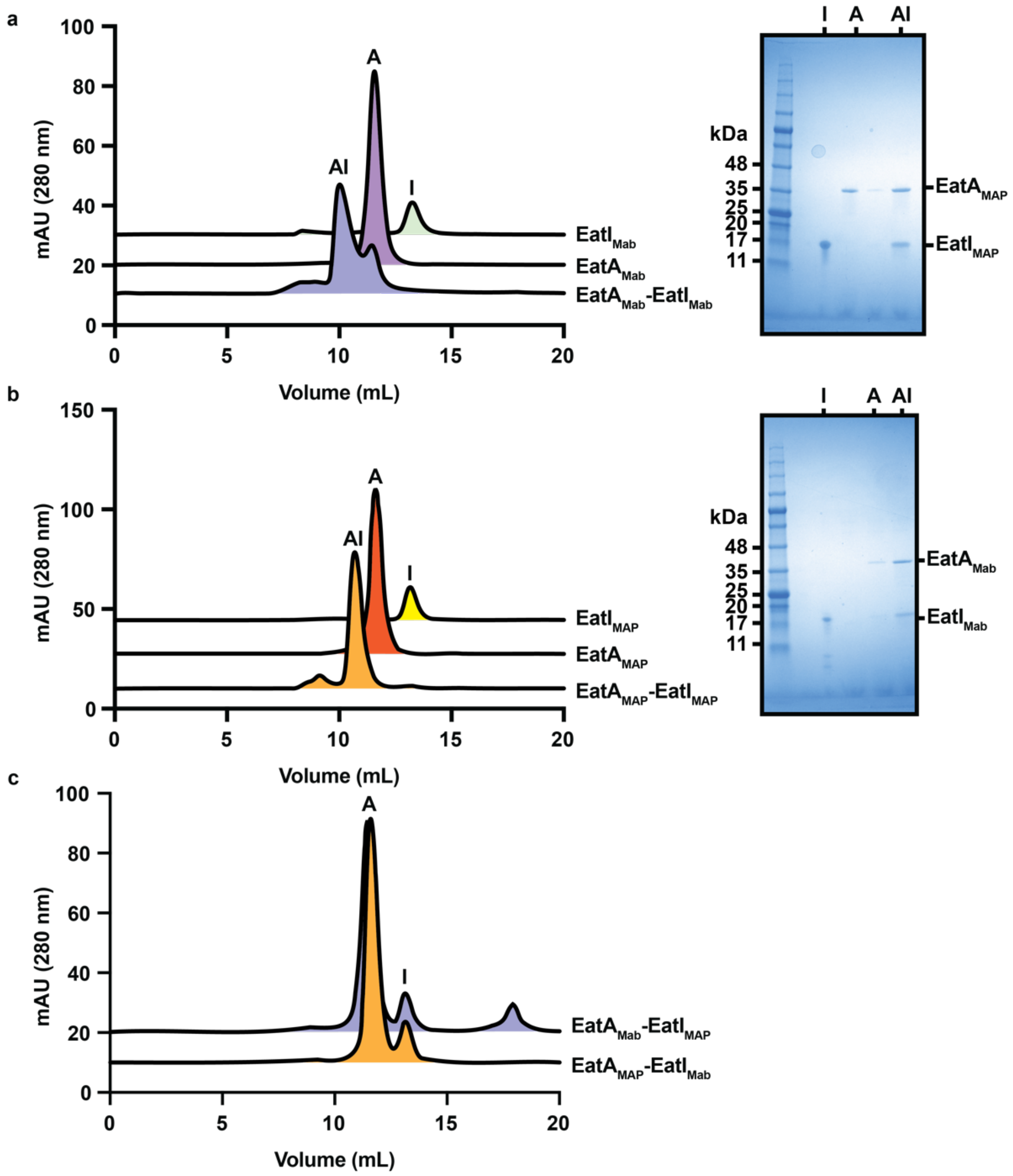


Recombinantly produced EatA GH183 domains and EatI proteins from *M. abscessus* (**a)** and *M. avium* subsp. *paratuberculosis* (**b**) were incubated alone or in combination and then analysed by size-exclusion chromatography on a Superdex 75 Increase 10/300 GL column. To identify peak fractions of the isolated proteins, higher molecular weight complexes were collected and analysed by SDS-PAGE (right). **c,** Non-cognate pairs of EatA and EatI were analysed similarly, demonstrating a lack of complex formation. The low molecular weight peak at 17.5 mL in EatI$_{Mab}$-EatA$_{MAP}$ is a contaminating protein.

**Extended Data Fig. 4. EatA enzymes are inhibited by cognate EatI proteins**

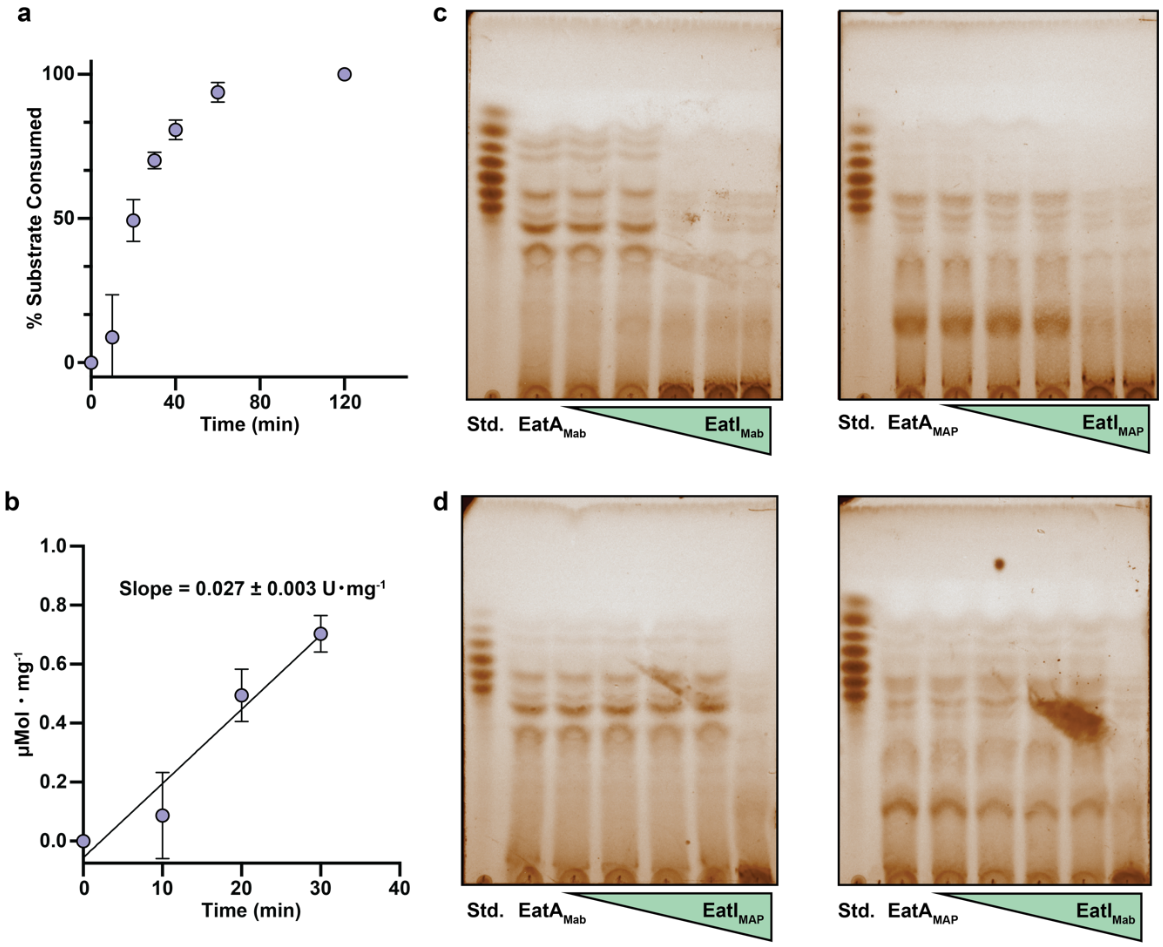


**a,** Substrate peaks from Fig. 2a were integrated and the percentage of substrate consumption relative to the amount of substrate at 0 min is reported. Error bars represent the standard deviation of three biological replicates. **b,** The linear portion of the curve in **a** was converted to μMol substrate conversion by comparing to standards of known concentration. A linear regression was computed in Graphpad Prism 10.4.1. Error bars represent the standard deviation of three biological replicates. $EatA_{Mab}$ and $EatA_{MAP}$ were incubated with their cognate (**c**) or non-cognate (**d**) EatI before initiating reactions with 1 mg・$mL^{-1}$ arabinogalactan from *M. smegmatis* $mc^2155$. Reactions were carried out at 0:1, 0.25:1, 0.5:1, 1:1, 2:1 and 2:0 EatI:EatA molar ratios for 30 min at 37 ℃ before being stopped by boiling, separated by TLC two times in a solvent system of 2:1:1 butanol:acetic acid:water and visualised by staining with orcinol and charring. Representative data from two biological replicates are presented. TLCs have been false-coloured in ImageLab 6.1 after black-and-white imaging.

**Extended Data Fig. 5. Structural analysis of substrate and immunity interaction interfaces in EatA proteins.**

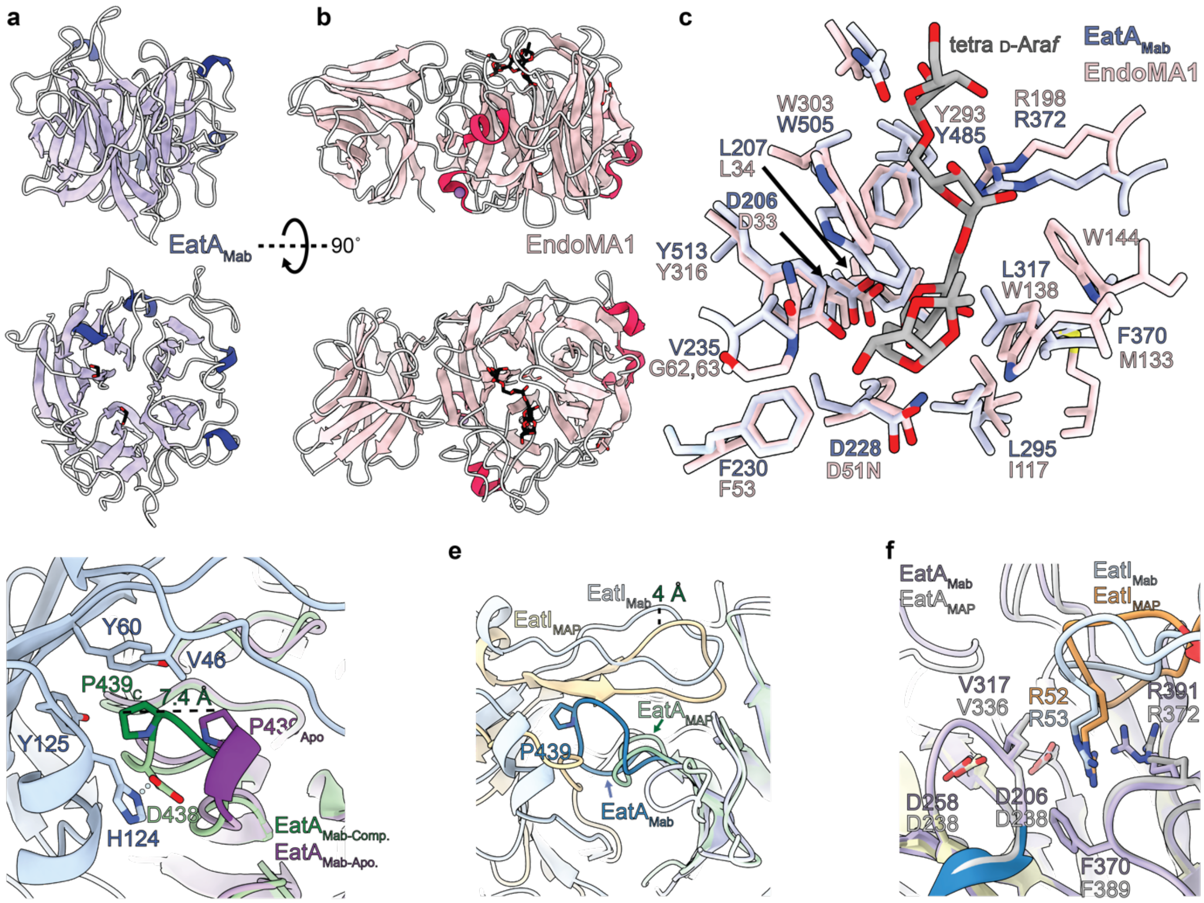


Aligned (RMSD = 5.48 Å) top and side views of $EatA_{Mab}$ (**a**) and EndoMA1 (**b;** PDB:8iC1). **c,** Key binding and catalytic ($EatA_{Mab}$ - D206, D228) residues are conserved between EndoMA1 and $EatA_{Mab}$. **d,** Binding of $EatI_{Mab}$ triggers an unwinding of a small helical turn (purple in $EatA_{Mab\text{-}Apo}$; green in $EatA_{Mab\text{-}Complex}$), moving P439 7.4 Å towards $EatI_{Mab}$. **e,** When aligned, the EatA-$EatI_{Mab}$ and EatA-$EatI_{MAP}$ structures have an overall RMSD of 2.6 Å. A key difference is the interface loop containing P439 in $EatA_{Mab,}$ which is substantially truncated in $EatA_{MAP}$. **f,** A conserved EatI Arg residue is coordinated in equivalent positions in the active sites in the EatA-$EatI_{Mab}$ and EatA-$EatI_{MAP}$ complexes.

**Extended Data Fig 6. An *M. abscessus* ATCC 1977 Δ*eccC4* mutant no longer secretes EsxU.**

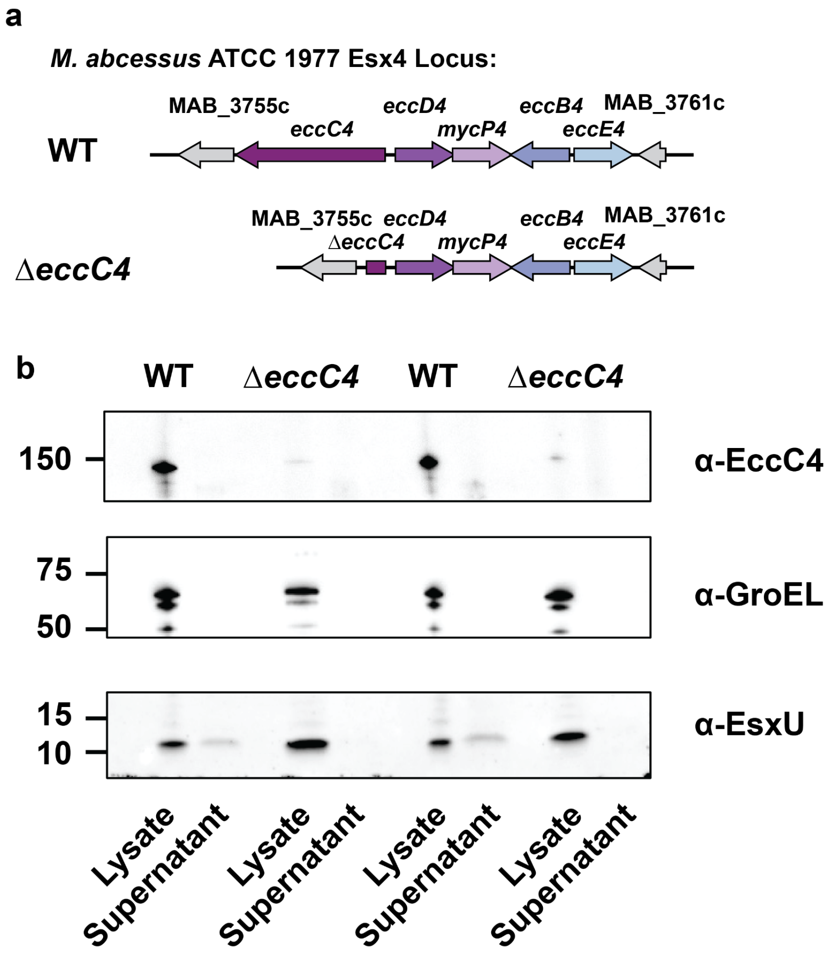


**a,** The *M. abscessus* ATCC 19977 *esx-4* locus in the wild-type and Δ*eccC4* mutant background. **b,** This mutation leads to a loss of function for ESX-4 as evidenced by the loss of EsxU secretion in the mutant culture supernatant. Left and right are biological replicates.

**Extended Data Table 1. Crystallography statistics.**

| | EatAMab | EatA-EatIMab | EatA-EatIMAP |
|---|---|---|---|
| **PDB Code:** | 9MXM | 9MYT | 9MZ5 |
| **Collection Statistics:** | | | |
| Wavelength | 0.9795 Å | 0.96863 Å | 0.6199 Å |
| Resolution range | 51.97 - 2.06 (2.1 - 2.06) | 89.917 - 1.702 (1.732 - 1.702) | 77.009 - 1.650 (1.792 - 1.650) |
| Space group | C 2 2 21 | P 1 21 1 | P 21 21 21 |
| Unit cell | 87.96 94.79 175.64 90 90 90 | 51.584 99.521 92.092 90 102.48 90 | 67.26 101.469 118.264 90 90 90 |
| Total reflections | 307785 | 575884 | 773467 |
| Unique reflections | 45733 | 89875 | 68287 |
| Multiplicity | 6.7 (7.1) | 6.4 (4.9) | 11.3 (6.6) |
| Completeness (%) | 100 (100) | 90.6 (49.2) | 88.5 (47.7) |
| Mean I/sigma(I) | 5.5 (0.8) | 6.3 (0.5) | 10.9 (1.6) |
| R-merge | 0.224 (2.177) | 0.137 (2.223) | 0.140 (1.078) |
| R-meas | 0.244(2.35) | 0.150 (2.482) | 0.146 (1.167) |
| R-pim | 0.095 (0.88) | 0.059 (1.069) | 0.041 (0.432) |
| CC1/2 | 0.989 (0.312) | 0.989 (0.37) | 0.998 (0.717) |
| **Refinement Statistics** | | | |
| Reflections used in refinement | 45680 | 89289 | 68264 |
| Reflections used for R-free | 2325 | 4238 | 3508 |
| R-work | 0.1922 | 0.1574 | 0.1601 |
| R-free | 0.2328 | 0.1973 | 0.1942 |
| Number of non-hydrogen atoms | 5481 | 8415 | 8053 |
| macromolecules | 5051 | 7309 | 6966 |
| ligands | 41 | 93 | 32 |
| solvent | 389 | 1013 | 1055 |
| Protein residues | 685 | 976 | 935 |
| RMS(bonds) | 0.014 | 0.008 | 0.010 |
| RMS(angles) | 1.22 | 0.93 | 1.09 |
| Ramachandran favored (%) | 95.74 | 97.42 | 97.51 |
| Ramachandran allowed (%) | 4.26 | 2.58 | 2.38 |
| Ramachandran outliers (%) | 0 | 0 | 0.11 |
| Rotamer outliers (%) | 0.56 | 0.38 | 0.53 |
| Clashscore | 5.84 | 3.09 | 5.79 |
| Average B-factor | 44.29 | 30.87 | 21.45 |
| macromolecules | 44.09 | 29.70 | 20.27 |
| ligands | 58.02 | 39.29 | 29.43 |
| solvent | 45.53 | 38.57 | 28.97 |

**Extended Data Table 2. Primers used in this study.**

| Construct | Oligonucleotides for cloning | |
|---|---|---|
| pqe70-*eatI*-*tapA1*-*tapA2*-TS-*eatA*-His$_6$ | backbone_F | catcaccatcaccatcactaagct |
| | backbone_R | catgcttaatttctcctctttaatgaattctgtgtg |
| | A1_F | atttcacacagaattcattaaagaggagaaattaagcatggcctgcaacgc atccg |
| | A1_R | ctcctgatcccggtcagaatctatatcttgggaatccagccggaac |
| | A2_F | gctggattcccaagatatagattctgaccgggatcaggagg |
| | A2_R | tgtggatgactccaagcactcatattctgtgggccggagg |
| | A3_F | cctccggcccacagaatatgagtgcttggagtcatccacaattc |
| | A3_R | tacatattctgtgggccggactacttctcaaattgtggatgagaccag |
| | A4_F | atccacaatttgagaagtagtccggcccacagaatatgtaggc |
| | A4_R | caagctcagctaattaagcttagtgatggtgatggtgatgttgatgtgggtttact tcgatcaattggg |
| pET28aEatA$_{Mab}$ D230A | D230AF | ggtttttggtgcgagcttcagcac |
| | D230AR | gctaccagtttttgcc |
| pMYBAD-*eat* | D230AF | tgtctttggagcgtctttcagcac |
| | D230AR | gcgacaagcttcttac |
| pUT18 | | ttgcatgcctgcaggtcg |
| pUT18_*tapA1* | | tgaccatgattacgccaagcagtgatgaacaggagctac |
| | | gtcgacctgcaggcatgcaagcggaacctctggaattg |
| pUT18_*tapA2* | | tgaccatgattacgccaagcggattgaacatagatcttgatcgtg |
| | | gtcgacctgcaggcatgcaacatattctgtgggccggag |
| pUT18_*eatA* | | tgaccatgattacgccaagcccgacgaaatcagaaattc |
| | | gtcgacctgcaggcatgcaattgatgtgggtttacttc |
| pUT18_*eatA*_N-terminus | | tgaccatgattacgccaagcccgacgaaatcagaaattcttaattg |
| | | gtcgacctgcaggcatgcaatcgatggccactgaagtc |
| pUT18_*eatA*_C-terminus | | tgaccatgattacgccaagcctgaagcccggcgaatctc |
| | | gtcgacctgcaggcatgcaattgatgtgggtttacttcgatcaattg |
| pT25 | | gtacctaagtaactaagaattcggc |
| | | ccggg gatcctctagagtc |
| pT25_*tapA1* | | cgactctagaggatccccggagtgatgaacaggagctac |
| | | attcttagttacttaggtacgcggaacctctggaattg |
| pT25_*tapA2* | | cgactctagaggatccccggggattgaacatagatcttgatcgtg |
| | | attcttagttacttaggtaccatattctgtgggccggag |
| pT25_*eatA* | | cgactctagaggatccccggccgacgaaatcagaaattc |
| | | attcttagttacttaggtacttgatgtgggtttacttc |
| pT25_*eatA*_N-terminus | | cgactctagaggatccccggccgacgaaatcagaaattcttaattg |
| | | attcttagttacttaggtactcgatggccactgaagtc |
| pT25_*eatA*_C-terminus | | cgactctagaggatccccggctgaagcccggcgaatctc |
| | | attcttagttacttaggtacttgatgtgggtttacttcgatcaattg |

**Extended Data Table 3. Predicted T7 Toxin sub-types identified**

| Toxin C-terminal module classification | Subgroup | C-terminal module prediction | Number of sequences within dataset (%) | Annotation/description |
|---|---|---|---|---|
| A | - | DUF4185/GH183 | 22 (1.5) | DUF4185 domain-containing protein; PF13810, $\beta$_Fructosidase Clan; glycoside hydrolase family 183 displays endo-$\alpha$-1,5-D-arabinofuranase activity[1]. |
| B | B1 | TNT | 10 (0.7) | TNT domain-containing protein; PF14021, ADP-ribosyl Clan; acts as a $NAD^+$ glycohydrolase triggering cell death. Putative binding pocket of Q822, Y765 and R757 has been identified in the CpnT of *M. tuberculosis*[2]. |
| B | B2 | TNT | 7 (0.5) | TNT domain-containing protein; PF14021, ADP-ribosyl Clan; acts as a $NAD^+$ glycohydrolase triggering cell death. Putative binding pocket of Q822, Y765 and R757 has been identified in the CpnT of *M. tuberculosis*[2]. |
| B | B3 | TNT | 35 (2.4) | TNT domain-containing protein; PF14021, ADP-ribosyl Clan; acts as a $NAD^+$ glycohydrolase triggering cell death. Putative binding pocket of Q822, Y765 and R757 has been identified in the CpnT of *M. tuberculosis*[2]. |
| C | C1 | a/b hydrolase 8 family | 20 (1.4) | $\alpha/\beta$ hydrolase; PF06259, $\alpha/\beta$_hydrolase Clan; contains a predicted Ser-His-Asp catalytic triad. Family named after characteristic 8 $\beta$-sheets, family covers a diverse range of enzyme classes[3]. |
| C | C2 | a/b hydrolase 8 family | 48 (3.3) | $\alpha/\beta$ hydrolase; PF06259, $\alpha/\beta$_hydrolase Clan; contains a predicted Ser-His-Asp catalytic triad. Family named after characteristic 8 $\beta$-sheets, family covers a diverse range of enzyme classes[3]. |
| C | C3 | a/b hydrolase 8 family | 49 (3.4) | $\alpha/\beta$ hydrolase; PF06259, $\alpha/\beta$_hydrolase Clan; contains a predicted Ser-His-Asp catalytic triad. Family named after characteristic 8 $\beta$-sheets, family covers a diverse range of enzyme classes[3]. |
| C | C4 | a/b hydrolase 8 family | 64 (4.4) | $\alpha/\beta$ hydrolase; PF06259, $\alpha/\beta$_hydrolase Clan; contains a predicted Ser-His-Asp catalytic triad. Family named after characteristic 8 $\beta$-sheets, family covers a diverse range of enzyme classes[3]. |
| D | D1 | NlpC/P60 | 84 (5.8) | NlpC/P60 family protein; PF00877, Peptidase_CA Clan; demonstrated to be involved in turnover of PG and essential to cell division[4]. |
| D | D2 | NlpC/P60 | 22 (1.5) | NlpC/P60 family protein; PF00877, Peptidase_CA Clan; demonstrated to be involved in turnover of PG and essential to cell division[4]. |
| E | - | Tox-REase-5 | 30 (2.1) | Tox-REase-5 domain-containing protein; PF15648, PDDEXK Clan; polymorphic toxin associated with multiple secretion systems and bacterial groups, possesses a restriction endonuclease fold (D-[EQ]xK)[5]. |
| F | F1 | Tox-REase-7 | 9 (0.6) | Putative toxin; PF15649, PDDEXK Clan; polymorphic toxin associated with multiple secretion systems and bacterial groups, possesses a restriction endonuclease fold (D-[EQ]xK)[5]. |
| F | F2 | Tox-REase-7 | 17 (1.2) | Putative toxin; PF15649, PDDEXK Clan; polymorphic toxin associated with multiple secretion systems and bacterial groups, possesses a restriction endonuclease fold (D-[EQ]xK)[5]. |

| | | | | |
|---|---|---|---|---|
| G | G1 | Ntox15 | 15 (1.0) | Hypothetical protein; PF15604; polymorphic toxin associated with multiple secretion systems and predicted to display RNase activity[5]. |
| | G2 | | 2 (0.1) | |
| H | - | Ntox37 | 35 (2.4) | Polymorphic toxin type 37; PF15535; polymorphic toxin associated with multiple secretion systems, possesses an all-b fold and predicted to display RNase activity[5]. |
| I | - | Ntox44 | 3 (0.2) | Polymorphic toxin type 44; PF15607; polymorphic toxin associated with multiple secretion systems, possesses an all-a-helical fold and predicted to display RNase activity[5]. |
| J | J1 | Nuclease | 5 (0.3) | Colicin-DNase; PF12639, His-Me_finger Clan; endonuclease domain that targets the ribosome to inactivate protein synthesis[6]. |
| | J2 | | 30 (2.1) | DNA/RNA non-specific endonuclease; PF13930, His-Me_finger Clan; possesses His-Me_finger fold observed in the *S. aureus* EsaD T7SSb antibacterial nuclease[7]. |
| | J3 | | 7 (0.5) | HNH/ENDO VII superfamily / GH-E family nuclease; PF14410, His-Me_finger Clan; NucA-like nuclease associated with multiple polymorphic toxins[8]. |
| | J4 | | 53 (3.7) | HNH/ENDO VII superfamily / GH-E family nuclease; PF14410, His-Me_finger Clan; NucA-like nuclease associated with multiple polymorphic toxins[8]. |
| K | - | Ribose hydrolase | 21 (1.5) | Hypothetical protein; PF02267, Nribosyltransf Clan; domain associated with the synthesis of cyclic ADP from NAD[9]. |
| L | - | CdiA_C_tRNase | 2 (0.1) | Hypothetical protein; PF18664, PDDEXK Clan; possesses tRNase $\alpha/\beta$-folds associated with PD(D/E)xK nucleases. Homologous to the type II CdiA from *Burkholderia*[10]. |

| | | | | |
|---|---|---|---|---|
| M | - | FGE-sulfatase | 15 (1.0) | Hypothetical protein; PF03781, C_Lectin Clan; eukaryotic-associated domain involved in targeting and modification of sulfate esters, has been shown to act as an iron(II)-dependent oxidoreductase in mycobacteria[11]. |
| N | - | ADP-ribosyltransferase | 3 (0.2) | ADP-ribosyltransferase; PF01129, ADP-ribosyl Clan; domain involved in post-transcriptional modification of proteins through transfer of NAD, homologous toxicity mechanism observed in the Cholera and Pertussis toxins[12]. |
| O | - | Colicin D | 13 (0.9) | Colicin D; PF11429, Colicin_D_E5 Clan; displays tRNase activity through targeting the phosphodiester bonds between the stem and loop[13]. |
| P | - | Amidase 6 | 37 (2.6) | Amidase domain-containing protein; PF12671, Peptidase_CA Clan; putative amidase domain that may act in a hydrolytic mechanism[14]. |
| Q | - | pApp synthetase | 41 (2.8) | ATP 3' pyrophosphokinase; cd05399, RelA/SpoT-like nucleotidyltransferase domain; domain associated with alarmone synthetases involved in gene regulation and co-ordinated stress responses[15]. |

**Note—Table Extended Data Table 3 details 699 toxin proteins comprising 48.6% of the 1,439-protein dataset. 677 (47%) proteins were classified EstU (EstU1 – U33) because their C-termini could not be assigned a predictive function. The remaining 63 (4.4%) protein sequences were removed throughout the analysis pipeline due to NCBI errors or FlaGs' rejection of accessions.**

#### Extended Data References

1. Al-Jourani, O. *et al.* Identification of d-arabinan-degrading enzymes in mycobacteria. *Nat. Commun.* **14**, 2233 (2023).

2. Sun, J. *et al.* The Tuberculosis Necrotizing Toxin kills macrophages by hydrolyzing NAD. *Nat Struct Mol Biol* **22**, 672–678 (2015).

3. Nardini, M. & Dijkstra, B. W. α/β Hydrolase fold enzymes: the family keeps growing. *Curr. Opin. Struct. Biol.* **9**, 732–737 (1999).

4. Griffin, M. E., Klupt, S., Espinosa, J. & Hang, H. C. Peptidoglycan NlpC/P60 peptidases in bacterial physiology and host interactions. *Cell Chem. Biol.* **30**, 436–456 (2023).

5. Zhang, D., Souza, R. F. de, Anantharaman, V., Iyer, L. M. & Aravind, L. Polymorphic toxin systems: Comprehensive characterization of trafficking modes, processing, mechanisms of action, immunity and ecology using comparative genomics. *Biol Direct* **7**, 18–18 (2012).

6. Soelaiman, S., Jakes, K., Wu, N., Li, C. & Shoham, M. Crystal Structure of Colicin E3 Implications for Cell Entry and Ribosome Inactivation. *Mol. Cell* **8**, 1053–1062 (2001).

7. Cao, Z., Casabona, M. G., Kneuper, H., Chalmers, J. D. & Palmer, T. The type VII secretion system of Staphylococcus aureus secretes a nuclease toxin that targets competitor bacteria. *Nat Microbiol* **2**, 16183 (2016).

8. Zhang, D., Iyer, L. M. & Aravind, L. A novel immunity system for bacterial nucleic acid degrading toxins and its recruitment in various eukaryotic and DNA viral systems. *Nucleic Acids Res.* **39**, 4532–4552 (2011).

9. Prasad, G. S. *et al.* Crystal structure of Aplysia ADP ribosyl cyclase, a homologue of the bifunctional ectozyme CD38. *Nat. Struct. Biol.* **3**, 957–964 (1996).

10. Johnson, P. M. *et al.* Unraveling the essential role of CysK in CDI toxin activation. *Proc. Natl. Acad. Sci.* **113**, 9792–9797 (2016).

11. Seebeck, F. P. In Vitro Reconstitution of Mycobacterial Ergothioneine Biosynthesis. *J. Am. Chem. Soc.* **132**, 6632–6633 (2010).

12. Okazaki, I. J., Kim, H.-J. & Moss, J. Cloning and Characterization of a Novel Membrane-associated Lymphocyte NAD:Arginine ADP-ribosyltransferase*. *J. Biol. Chem.* **271**, 22052–22057 (1996).

13. Yajima, S. *et al.* Relation between tRNase activity and the structure of colicin D according to X-ray crystallography. *Biochem. Biophys. Res. Commun.* **322**, 966–973 (2004).

14. Dong, C. *et al.* Structural insights into the inhibition of type VI effector Tae3 by its immunity protein Tai3. *Biochem. J.* **454**, 59–68 (2013).

15. Chakraburtty, R., White, J., Takano, E. & Bibb, M. Cloning, characterization and disruption of a (p)ppGpp synthetase gene (relA) of Streptomyces coelicolor A3(2). *Mol. Microbiol.* **19**, 357–368 (1996).
